# Fast, multiplexable and highly efficient somatic gene deletions in adult mouse skeletal muscle fibers using AAV-CRISPR/Cas9

**DOI:** 10.1101/2023.02.20.529232

**Authors:** Marco Thürkauf, Shuo Lin, Filippo Oliveri, Dirk Grimm, Randall J. Platt, Markus A. Rüegg

**Affiliations:** Biozentrum, University of Basel, Basel, Switzerland; Department of Infectious Diseases/Virology, Section Viral Vector Technologies, Medical Faculty, Heidelberg University, Heidelberg, Germany; BioQuant, University of Heidelberg, Heidelberg, Germany; German Center for Infection Research (DZIF) and German Center for Cardiovascular Research (DZHK), Heidelberg, Germany; Department of Biosystems Science and Engineering (D-BSSE), ETH Zurich, Basel, Switzerland; Department of Chemistry, University of Basel, Basel, Switzerland

## Abstract

Molecular screens comparing different disease states to identify candidate genes rely on the availability of fast, reliable and multiplexable systems to interrogate genes of interest. CRISPR/Cas9- based reverse genetics is a promising method to eventually achieve this. However, such methods are sorely lacking for multi-nucleated muscle fibers, since highly efficient nuclei editing is a requisite to robustly inactive candidate genes. Here, we couple Cre-mediated skeletal muscle fiber-specific Cas9 expression with myotropic adeno-associated virus-mediated sgRNA delivery to establish a system for highly effective somatic gene mutation in mice. Using well-characterized genes, we show that local or systemic inactivation of these genes copy the phenotype of traditional gene-knockout mouse models. Thus, this proof-of-principle study establishes a method to unravel the function of individual genes or entire signaling pathways in adult skeletal muscle fibers without the cumbersome requirement of generating knockout mice.

## Introduction

With the advent of –omics technologies that allow to correlate molecular signatures with specific disease states of cells or tissues, there is an increasing need for methods to interrogate the function of genes and pathways. Traditionally, forward and reverse genetics using targeted mutagenesis in combination with transgenesis has been used. More recently, clustered regularly interspaced short palindromic repeats (CRISPR)-mediated genome editing has become the method of choice for gene engineering in many species and tissues (*1*).

When it comes to skeletal muscle tissue, studying gene function *in vivo* is particularly challenging. Skeletal muscle is one of the largest organs constituting up to 50% of the mammalian body mass (*2*). The size and the fact that muscle fibers, which are the functional contractile units of skeletal muscle, form a syncytium with hundreds of myonuclei in a common cytosol, represent a substantial challenge for somatic gene inactivation. Therefore, the method of choice for functional gene interrogation studies in muscle remains transgenic mice generated *via* the Cre-loxP system. However, generation of transgenic mice requires extensive breeding, making functional interrogation of multiple genes cumbersome and time consuming.

Effective methods for somatic gene perturbation would offer huge advantages for screening multiple muscle gene candidates. While RNA interference, which can silence a target gene by introducing short hairpin (sh) RNAs (*3*), can acutely silence gene expression in muscle fibers (*4, 5*), prolonged elimination of a gene product requires sustained, high expression of the shRNA. The introduction of viruses, in particular adeno-associated viruses (AAV), as vehicles for delivering shRNAs, opened the possibility of systemic administration (*6*). However, due to the lack of tissue-specific control of shRNA expression, gene silencing occurs in all transduced cells. While next-generation AAV capsids with designed tropism towards skeletal muscle tissue (*7–9*) may improve the off-tissue targeting, all of them also target myocytes in the heart. Another challenge for somatic gene targeting of muscle fibers is the overall heterogeneity of the tissue. Almost half of the nuclei in skeletal muscle derive from non-fiber cells, such as muscle stem cells (MuSC), endothelial cells, fibro-adipogenic precursors (FAPs), Schwann cells or tenocytes (*10*) and perturbation of their function often affects muscle fibers as well. Therefore, for rapid functional gene interrogation in skeletal muscle fibers, an efficient, multiplexable and muscle fiber-specific gene editing approach is sorely needed.

Here we establish a versatile tool for local and systemic skeletal muscle fiber-specific gene knockout. This tool couples the advantages of CRISPR with newly developed, highly efficacious, AAV9- derived viral capsids by using (i) mice engineered to constitutively or inducibly express Cas9 in skeletal muscle fibers and (ii) delivering single guide (sg) RNAs with the myotropic AAVMYO (*7*). By targeting key genes, we demonstrate that this system is capable of potently altering signaling pathways, destroying neuromuscular junctions and stimulating muscle hypertrophy without needing to generate germline gene-of-interest deletions.

## Results

### Constitutive expression of Cas9 in skeletal muscle fibers

To express Cas9 at high levels in skeletal muscle fibers, we crossed Cre-dependent Rosa26^Cas9-EGFP^ knockin mice (*11*) with mice expressing Cre recombinase constitutively (scheme in Fig. 1A) or upon tamoxifen injection (scheme in Fig. S2A, S2B) in skeletal muscle fibers (*12, 13*). The resulting transgenic mice were called Cas9mKI and iCas9mKI mice, respectively. Expression of Cas9 in Cas9mKI mice was confirmed by immunohistochemistry for GFP (Fig. 1B). By Western blot analysis, Cas9 expression was detected in all muscles tested but not in heart or liver (Fig. 1C). Most importantly, skeletal muscle mass and function (Fig. 1D-I) as well as fiber-type composition and neuromuscular junction (NMJ) structure (Fig. S1) of adult Cas9mKI mice were indistinguishable from control mice. Similarly, fourteen days after tamoxifen injection, Cas9 was high in adult skeletal muscle but not detected in heart or liver of iCas9mKI mice (Fig. S2). Together, these data confirm the strong and tissue-restricted expression of Cas9 in skeletal muscle fibers of Cas9mKI and iCas9mKI mice.

**Figure 1:**
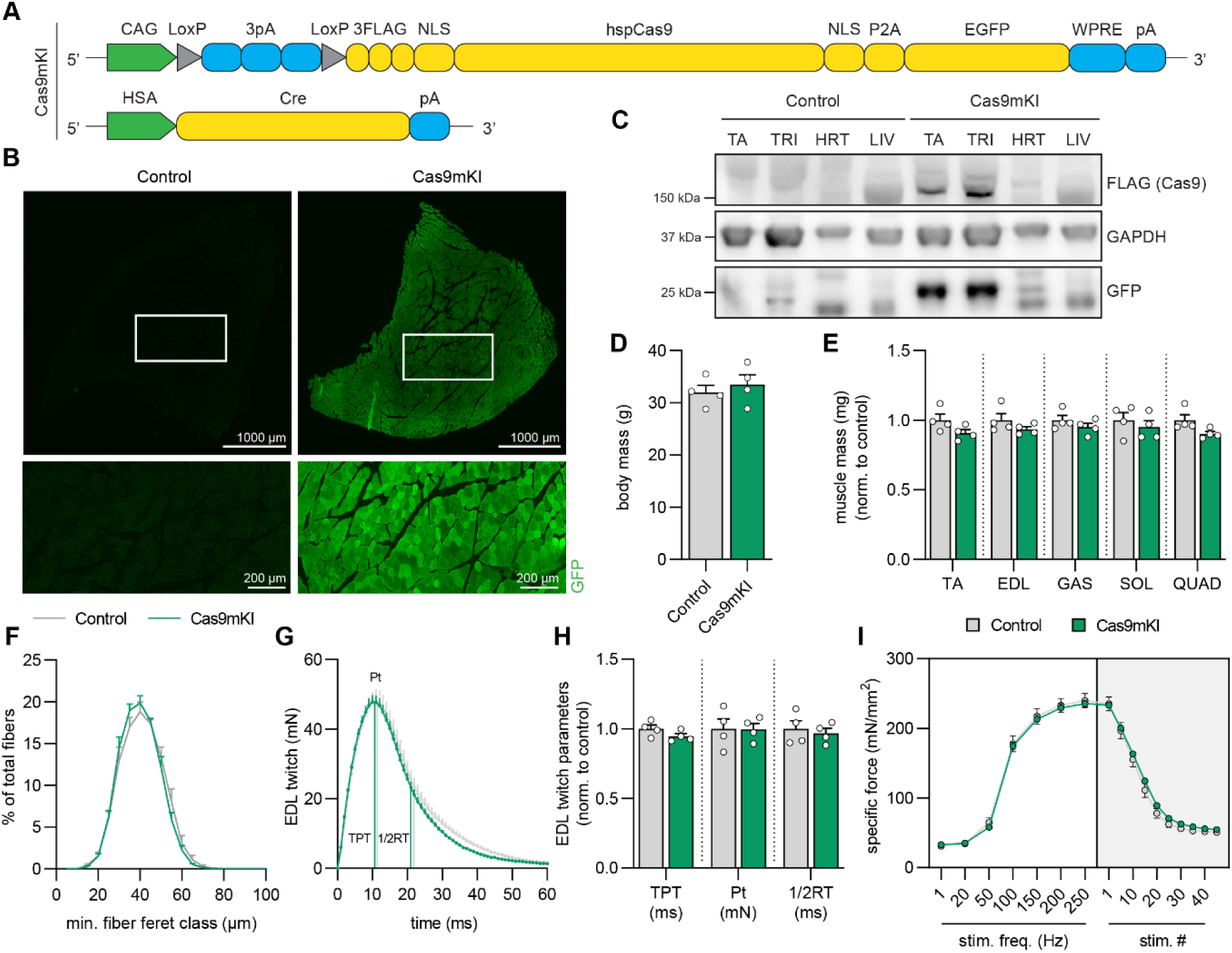
Validation of Cas9mKI mice. (A) Schematic of the Cas9mKI mouse model. Abbreviations are: CAG: cytomegalovirus (CMV) enhancer fused to the chicken beta-actin promoter; LoxP: locus of X-over P1; pA: polyadenylation signal; FLAG: FLAG-tag; NLS: nuclear localization signal; hspCas9: humanized *Streptococcus pyogenes* Cas9; P2A: 2A self-cleaving peptide; EGFP: enhanced green fluorescent protein; WPRE: woodchuck hepatitis virus post-transcriptional regulatory element; HSA: human α- skeletal actin; Cre: Cre recombinase. (B) Cross-sections of *tibialis anterior* (TA) muscle stained for EGFP (green) in control and Cas9mKI mice. (C) Western blot analysis of lysates from TA, *triceps brachii* (TRI), heart (HRT) and liver (LIV) of control and Cas9mKI mice using antibodies against the FLAG-tag, GFP or glyceraldehyde-3-phosphate dehydrogenase (GAPDH). Only TA and TRI muscles of Cas9mKI mice are positive for the FLAG-tag and GFP; no expression was detected in HRT or LIV. (D) Body mass of 19-week-old control and Cas9mKI mice. (E) Relative mass of *soleus* (SOL), *extensor digitorum longus* (EDL), TA, *gastrocnemius* (GAS) and *quadriceps* (QUAD) muscles from control and Cas9mKI mice. (F) Minimal fiber feret distribution of muscle fibers from TA of control and Cas9mKI mice. (G) Ex-vivo twitch response of isolated EDL muscle from Cas9mKI and control mice. Peak twitch (Pt), time-to-peak twitch (TPT) and half-relaxation time (1/2RT) are indicated. (H) Quantification of ex-vivo twitch response parameters (TPT, Pt, 1/2RT) of isolated EDL muscle from Cas9mKI and control mice. (I) Force-frequency curve (left) and fatigue response to multiple stimulations (right) of EDL muscle from control and Cas9mKI mice. Data are means ± SEM. N = 4 (mice). None of the data are significantly different between control and Cas9mKI mice (P > 0.05) using unpaired t-test.

### Robust *in vivo* gene editing using local AAV9-mediated sgRNA delivery into Cas9mKI mice

To test whether high Cas9 expression would allow gene perturbation in skeletal muscle to an extent required to lower protein levels, we selected *Prkca*, which codes for protein kinase Cα (PKCα). We selected PKCα based on a combination of our experience characterizing PKCα as an mTORC2 target in the brain (*14*), the availability of antibodies for Western blot analyses and because CRISPR has been used to successfully eliminate PKCα in the retina (*15*). As low PKCα levels, due to loss of mTORC2, do not affect skeletal muscle (*16*), we could determine the effectiveness of the system independent of secondary effects by the loss of PKCα. Beside the published sgRNA (called sgPKCα-1), we tested initially an additional sgRNA (sgPKCα-2) and included a non-targeting sgRNA (sgNT). Cultured C2C12 myoblasts were transfected with a plasmid encoding the U6 promoter-driven sgRNA followed by an EFS promoter-driven Cas9, the P2A self-cleavage peptide and puromycin N-acetyltransferase, which confers puromycin resistance to transfected cells (Fig. S3A). After puromycin-selection, C2C12 myoblasts were differentiated into myotubes for five days. All selected cells expressed Cas9 and those co-expressing sgPKCα-1 or sgPKCα-2, but not sgNT, showed strongly reduced levels of PKCα (Fig. S3B, C). As sgPKCα-1 and sgPKCα-2 showed similar efficiency, we selected the published sgPKCα-1 (*15*) for further characterization. To quantify the number of <u>in</u>sertions and <u>del</u>etion<u>s</u> of bases (indels), we sequenced genomic DNA in the region targeted by sgPKCα-1 and used the method of “<u>T</u>racking of Indels by <u>De</u>composition” (TIDE). The total DNA-editing efficiency for sgPKCα-1 was 62.4 ± 1.6% (Fig. S3D), with the majority of deletions lacking 1 base pair (−1 bp) followed by insertions of +1 bp (Fig. S3E).

Based on the high efficiency of sgPKCα-1 in editing *Prkca* and lowering the amount of PKCα in cultured C2C12 cells, we next injected AAV9 expressing three copies of sgPKCα-1 and co-expressing tdTomato under the CMV promoter (scheme in Fig. 2A) into *tibialis anterior* (TA) muscle of adult Cas9mKI mice. Additionally, we co-injected neuraminidase, which has been shown to improve AAV9 transduction of skeletal muscle (*17, 18*). We used a non-targeting sgRNA (sgNT) as a control. Six weeks after injection of either AAV9-sgNT or AAV9-sgPKCα-1 (3 x 10^11^ vg) into TA muscle of Cas9mKI mice, several tissues were analyzed. Transduction efficiency was monitored by staining for tdTomato in TA muscle cross-sections (Fig. 2B) and by measuring AAV genome copy numbers per nucleus in different tissues (Fig. 2C). Expression of tdTomato was quite homogenous (Fig. 2B) and transduction rates reached 99 ± 9.6 vg/nucleus in TA (Fig. 2C). Virus leakage into the blood stream resulted in strong liver (117.8 ± 20.0 vg/nucleus) and weak heart (19 ± 3.3 vg/nucleus) transduction (Fig. 2C). To determine genome editing efficiency, we again used TIDE in the targeted *Prkca* locus on DNA isolated from AAV9- sgNT or AAV9-sgPKCα-1-transduced TA muscle (Fig. 2D). The background editing signal in AAV9-sgNT-transduced muscle was 1.4 ± 0.5%, while the experimental muscle reached 20.3 ± 1.0% editing (Fig. 2D, E). As a consequence of CRISPR/Cas9-mediated DNA editing, PKCα protein was strongly diminished in AAV9-sgPKCα-1-injected compared to AAV9-sgNT-injected TA muscle (Fig. 2F, G). The low amount of PKCα still detected in AAV9-sgPKCα-1-transduced TA muscle may also derive from other muscle-resident cells that express *Prkca* transcripts (*10*). Together, these data show that neuraminidase treatment coupled with AAV9-mediated sgRNA delivery markedly reduced PKCα protein in the targeted skeletal muscle of Cas9mKI mice.

**Figure 2:**
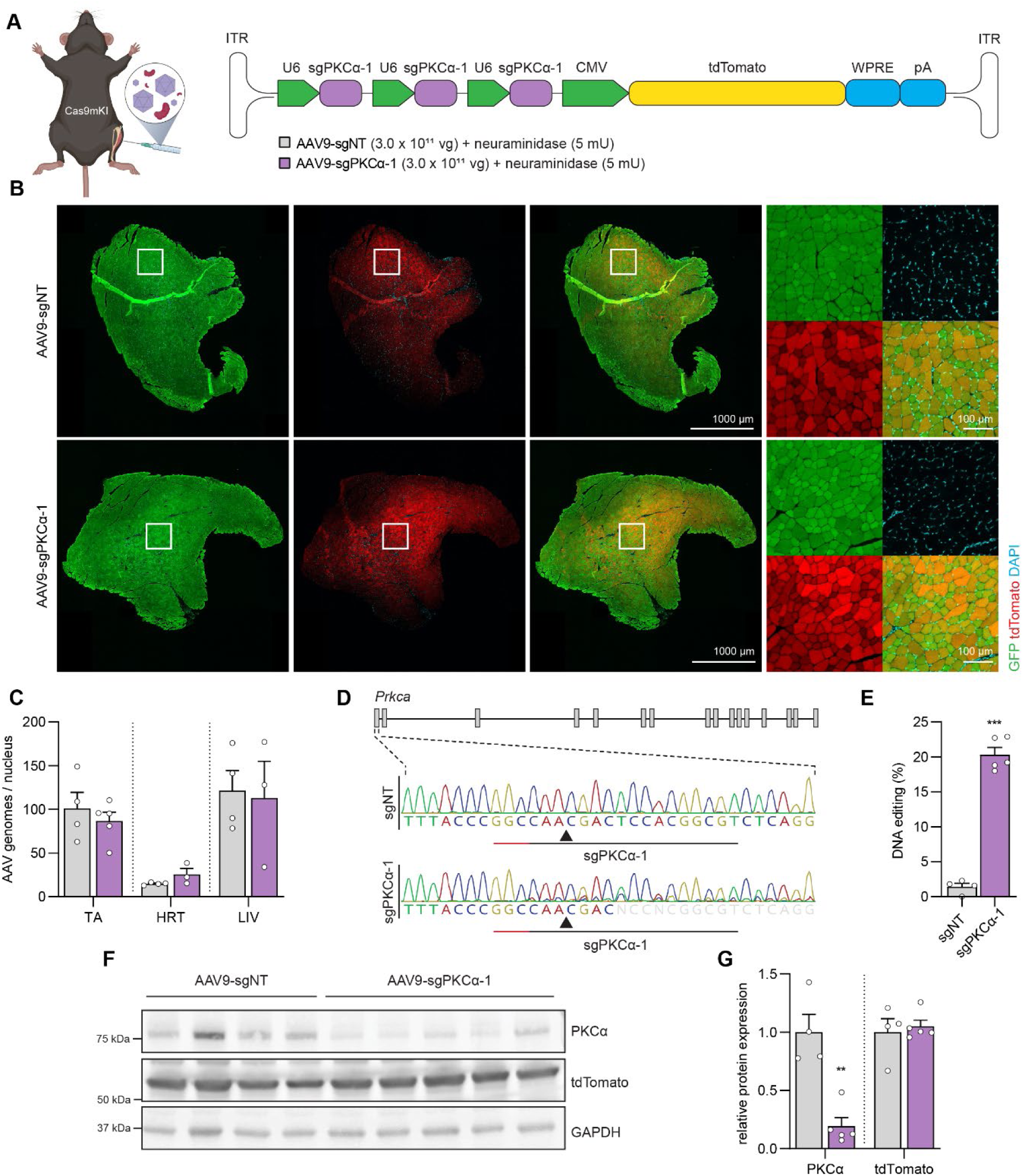
Knock out of *Prkca* in TA muscle upon AAV9/neuraminidase-mediated delivery of sgRNA to Cas9mKI mice. (A) Schematic illustration of experimental procedure and targeting construct. For abbreviations, see legend to Figure 1. (B) Cross-section of *tibialis anterior* (TA) muscle stained for Cas9/GFP (green), tdTomato (red) and DAPI (blue) 6 weeks after injection of neuraminidase plus AAV9- sgNT or AAV9-sgPKCα-1. (C) Quantification of AAV genomes per nucleus in TA muscle, heart (HRT) and liver (LIV) in AAV9-sgNT (grey) and AAV9-sgPKCα-1-injected (purple) mice. (D) Illustration of the *Prkca* gene and representative Sanger sequencing chromatograms at the sgPKCα-1 target site (underlined) of AAV9-sgNT and AAV9-sgPKCα-1-injected TA muscle. Note that sequencing becomes ambiguous in sgPKCα-1-expressing mice, indicative of genome editing. (E) Total INDEL formation analysis by TIDE of TA muscle injected with AAV9-sgNT and AAV9-sgPKCα-1. (F) Western blot analysis and (G) quantification of PKCα and tdTomato expression in TA muscle 6 weeks post-injection with AAV9-sgNT (grey) or AAV9-sgPKCα-1 (purple). Data are means ± SEM. N = 4 (sgNT) and 5 (sgPKCα-1) mice. Statistical analysis used unpaired t-test. *P < 0.05, **P< 0.01, ***P < 0.001 .

### Improved editing efficiency with AAVMYO for local sgRNA delivery into Cas9mKI mice

To further improve DNA editing and facilitate systemic applications (which is not possible when injecting neuraminidase), we next tested a peptide-displaying AAV9 capsid variant, called AAVMYO, with superior skeletal muscle fiber tropism (*7*). We compared the efficiency of AAVMYO-sgPKCα-1 with AAV9-sgPKCα-1 by injecting 3 x 10^11^ vg of each, or a PBS control into TA muscle (Fig. 3A). Six weeks post-injection, tdTomato expression was visibly higher in TA muscle as well as the nearby *extensor digitorum longus* (EDL) and *gastrocnemius* (GAS) muscles of AAVMYO-injected mice than AAV9- injected muscles (Fig. 3B). Cross sections from TA (Fig. 3B), EDL and GAS muscles (Fig. S4A) as well as Western blot quantification from TA muscle (Fig. 3C-D) further confirmed higher tdTomato expression in AAVMYO- than AAV9-injected muscles. Average transduction by AAVMYO, judged by AAV genomes/nucleus, was at least 2.5 times higher than by AAV9 for all muscles, including the heart, while transduction of the liver was markedly lower (Fig. S4B). The superior transduction efficiency of AAVMYO over AAV9 upon intramuscular injection is in line with previous observations upon systemic administration of AAVMYO and AAV9 (*7*). As a consequence of the more efficient transduction, the amount of PKCα was also strongly diminished in AAVMYO-sgPKCα-1-transduced TA (Fig. 3C, D), EDL (Fig. S4C, D) and GAS (Fig. S4F, G) muscles compared to AAV9-sgPKCα-1. TIDE analysis showed a higher total editing efficiency of the sgPKCα-1-targeted locus by AAVMYO-sgPKCα-1 (22.4 ± 1.4%) than AAV9- sgPKCα-1 (17.2 ± 0.9%) in TA muscle (Fig. 3E). Compared to the intramuscular injection of AAVMYO into TA muscle, gene editing and knockdown efficiencies remained very similar in the adjacent EDL and GAS muscles, while both values dropped with AAV9 (Fig. S4E, H).

**Figure 3:**
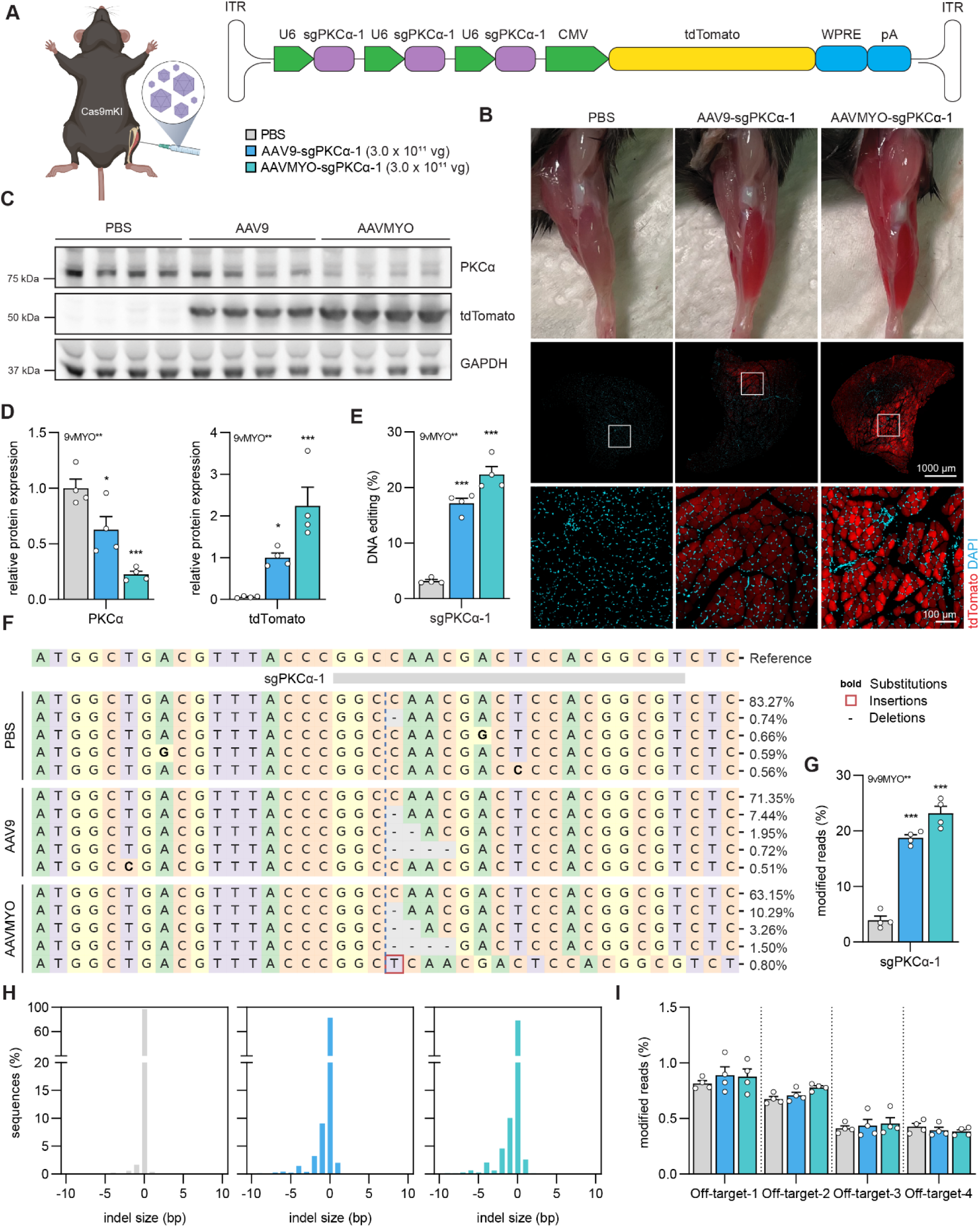
AAVMYO-mediated sgPKCα-1 delivery into TA muscle results in strong reduction of PKCα. (A) Schematic presentation of the experimental procedure. For abbreviations, see legend to Figure 2. (B) Representative images of the injected hindlimb and of cross-sections of *tibialis anterior* (TA) muscle stained for tdTomato (red) and DAPI (blue), 6 weeks post-intramuscular injection of PBS or AAVs (3.0 x 10^11^ vg) into Cas9mKI mice. (C) Western blot analysis and quantification (D) for PKCα and tdTomato in TA muscle lysates of Cas9mKI mice injected with PBS (grey), AAV9-sgPKCα-1 (light blue) or AAVMYO- sgPKCα-1 (cyan) 6 weeks post-injection. (E) Total INDEL formation analysis by TIDE. (F) Representative sequence frequency table of reads using DNA isolated from TA under the different conditions covering the sgPKCα-1 target region. (G) Relative number of modified reads under the different conditions in the sgPKCα-1 target region. (H) INDEL size histogram indicating mutation distribution at the sgPKCα-1 target region in TA muscle of Cas9mKI mice. Conditions are injection of PBS (light grey, left), AAV9- sgPKCα-1 (light blue, middle) or AAVMYO-sgPKCα-1 (cyan, right). (I) Total amount of mutated reads of amplicons covering the top four predicted off-target loci in the different experimental paradigms. There is no difference in the modified reads compared to PBS injection. Data are means ± SEM. N = 4 mice for each condition. Statistical significance is based on one-way ANOVA with Fishers LSD post-hoc test. *P < 0.05, **P < 0.01, ***P < 0.001.

To more precisely map genome editing frequency in the genomic DNA surrounding the sgPKCα-1 target site, we performed next-generation sequencing (NGS) of TA muscle DNA (Fig. 3F and Table S1). The sum of all observed mutations with NGS was comparable to TIDE analysis; with average mutations of 18.7 ± 0.6% for AAV9 and 23.1 ± 1.3% for AAVMYO (Fig. 3G). Independent of the AAV capsid variant, the most frequent indels were short deletions (Fig. 3H). To test whether introduction of sgPKCα-1 caused off-target editing, we also sequenced the genome in the top four off-target sites as predicted by the CRISPR-design tool CRISPOR (*19*). No significant sequence alterations were detected at these loci (Fig. 3I).

**Table 1:**
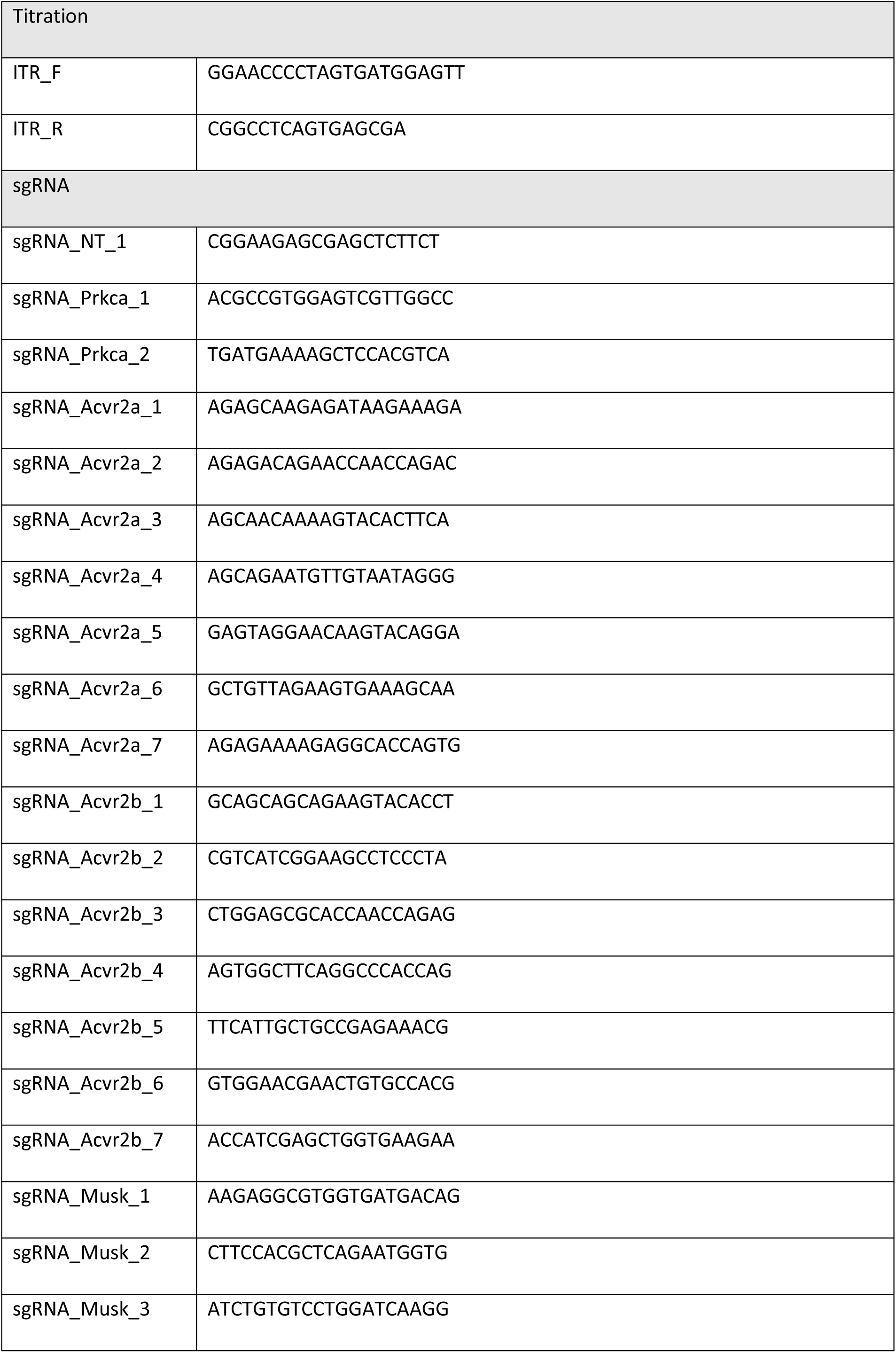
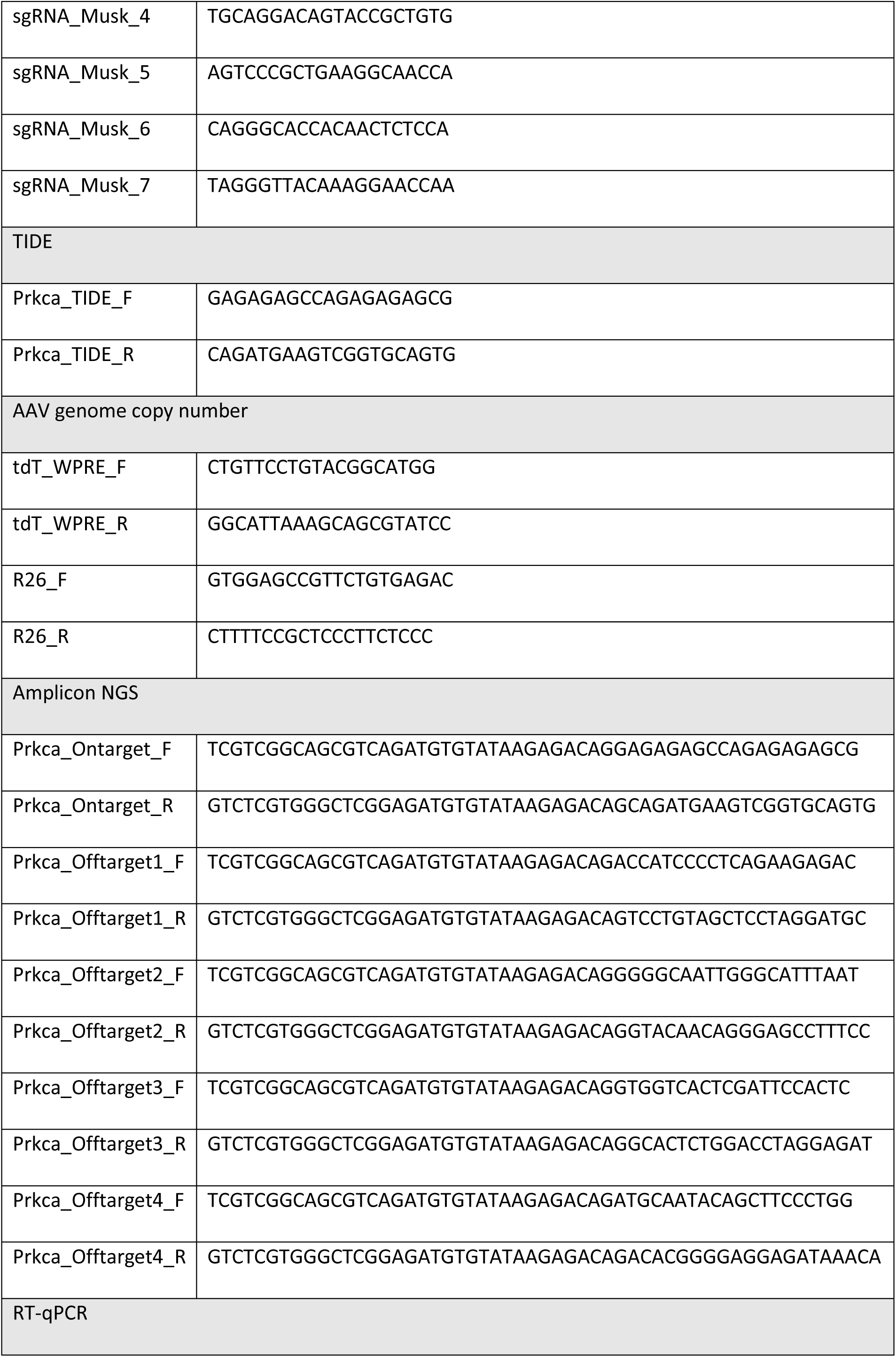
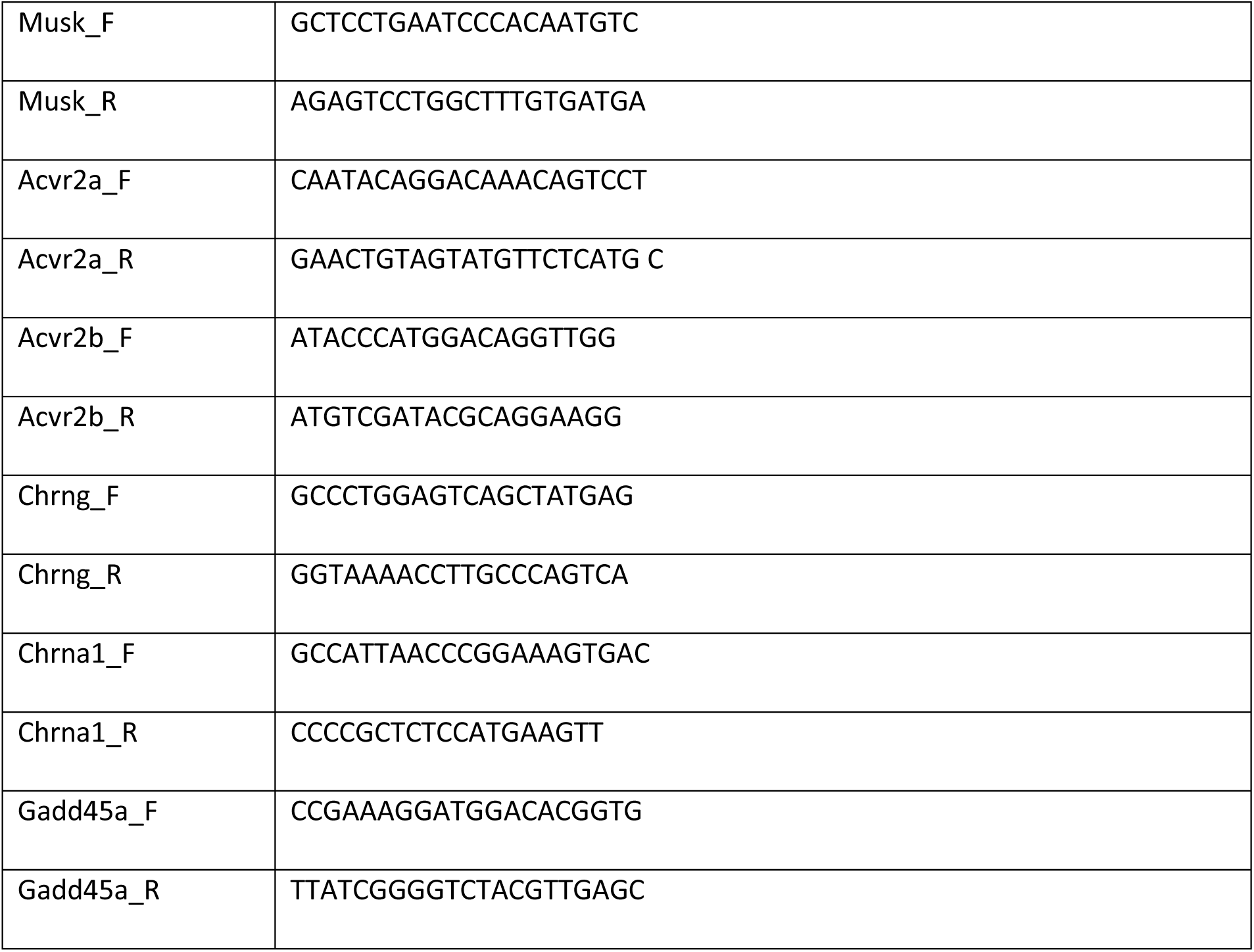
List of used primers and sgRNAs

Denervation and hence loss of muscle contraction has an immediate effect on gene expression in myonuclei and results after a few days in exuberant muscle atrophy. It indirectly also affects satellite cells and many other muscle-resident mononuclear cells. As somatic gene deletion may affect innervation, we wanted to assess whether the gene editing system would also work during acute denervation. To test this, we injected AAVMYO-sgPKCα-1 (3 x 10^11^ vg) or PBS (as a control) into TA muscle of Cas9mKI mice before unilateral sciatic nerve transection 6 weeks later and then analyzed muscle 14 days later. The denervation-induced loss of muscle mass was not different between PBS and AAVMYO-sgPKCα-1-injected mice (Fig. S5A). Importantly, denervation did not affect the overall PKCα knockdown efficiency or expression of the denervation marker HDAC4 (Fig. S5B, C). There was a slight decrease in the total percentage of genome editing by sgPKCα-1 (Fig. S5D), which is likely due to the increase in non-muscle fiber cells following denervation that do not express Cas9 (*20*). Together, our data show that AAVMYO-mediated sgRNA delivery into TA muscle induces robust and specific *in vivo* gene perturbation.

### AAVMYO supersedes AAV9 for systemic sgRNA delivery

To evaluate efficiency for systemic gene editing, we next injected 1 x 10^14^ vg/kg of AAV9-sgPKCα-1 or AAVMYO-sgPKCα-1 into the tail vein of 6-week-old Cas9mKI mice and collected tissues 6 weeks later (scheme Fig. 4A). Expression of tdTomato was visually higher at autopsy and strikingly higher in cross-sections of multiple muscles in mice injected with AAVMYO than with AAV9 (Fig. 4B). Similar results were obtained for the heart (Fig. S6A). Numbers of viral genomes per nucleus were 4- to 6-fold higher in limb muscles (TA, EDL, SOL and TRI) and more than 13-fold higher in the diaphragm (DIA) with AAVMYO than AAV9 (Fig. 4C). In line with the high transduction efficiency, AAVMYO-sgPKCα-1 induced 2.2- to 7.6-fold higher DNA editing rates across different muscles than AAV9-sgPKCα-1 (Fig. 4D). The most striking difference was seen in DIA muscle, where AAVMYO-sgPKCα-1 induced 19.1 ± 0.9% DNA editing while AAV9-sgPKCα-1 induced only 2.5 ± 0.9%. Western blot analysis for tdTomato and PKCα confirmed the superior systemic transduction of muscle tissue by AAVMYO-sgPKCα-1, with higher tdTomato expression and stronger reduction in PKCα protein abundance than with AAV9-sgPKCα-1 (Fig. 4E-G).

**Figure 4:**
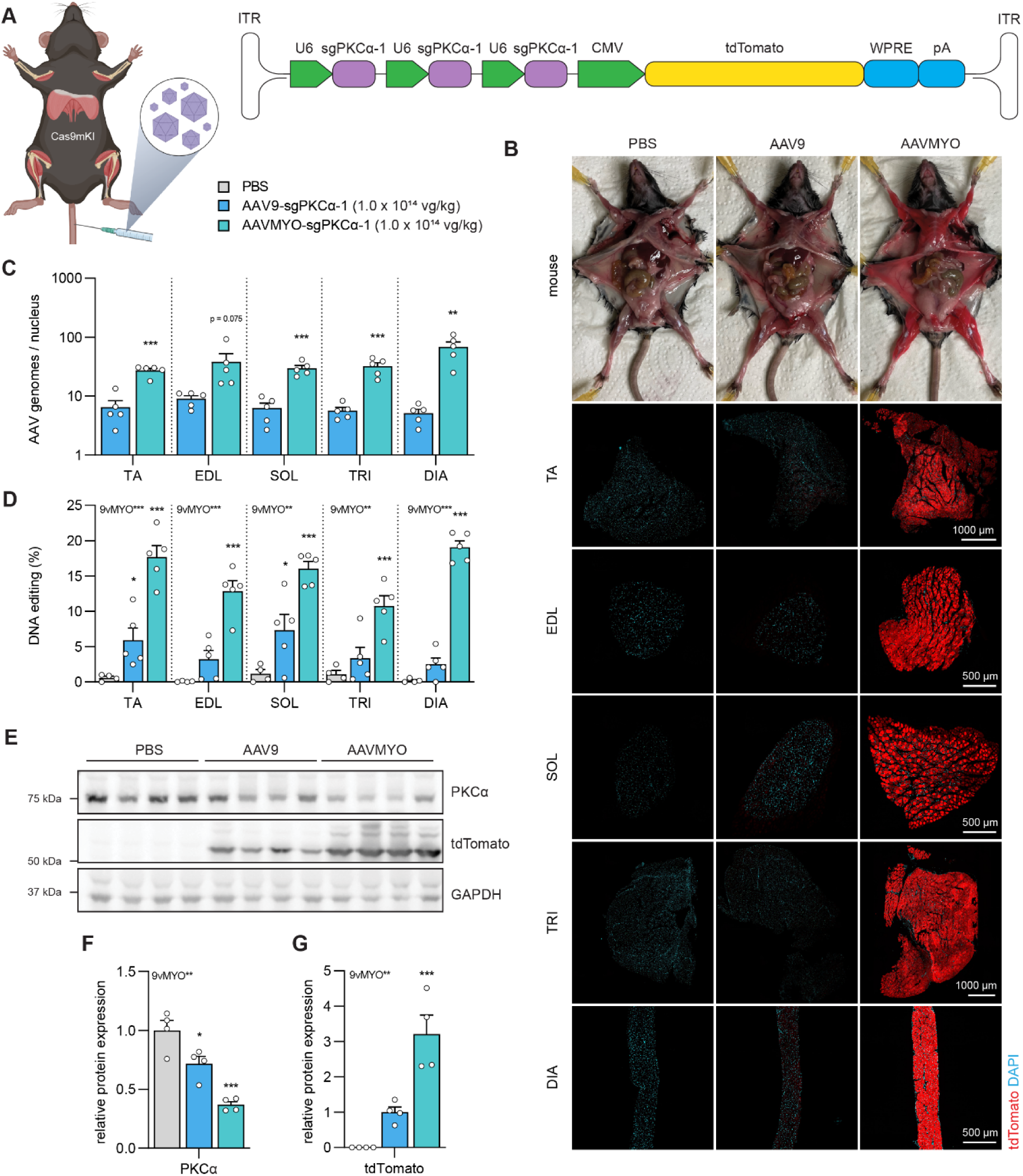
Systemic administration of AAVMYO-sgPKCα-1 *via* the tail vein into Cas9mKI mice reduces PKCα protein. (A) Schematic presentation of the experimental procedure. For abbreviations, see legend to Figure 2. (B) Representative images of dissected mice and cross-sections of *tibialis anterior* (TA), *extensor digitorum longus* (EDL), *soleus* (SOL), *triceps brachii* (TRI) or diaphragm (DIA) muscle stained for tdTomato (red) and DAPI (blue), 6 weeks post-intravenous injection of PBS or AAV (1.0 x 10^14^ vg/kg) into Cas9mKI mice. (C) Distribution of AAV in TA, EDL, SOL, TRI and DIA upon intravenous injection of AAV9-sgPKCα-1 (light blue) and AAVMYO-sgPKCα-1 (cyan) into Cas9mKI mice. (D) Total INDEL formation analysis by TIDE. (E) Western blot analysis and quantification (F, G) for PKCα (F) and tdTomato (G) in TA muscle of Cas9mKI mice injected with PBS (grey), AAV9-sgPKCα-1 (light blue) or AAVMYO-sgPKCα-1 (cyan). Data are means ± SEM. N = 4 - 5 mice. Significance was determined using one-way ANOVA with Fishers LSD post-hoc test (D, F, G) or unpaired t-test (C). *P < 0.05, **P < 0.01, ***P < 0.001.

We also tested systemic administration of additional capsid variants of AAVMYO, called AAVMYO2 and AAVMYO3, that were originally selected for their liver de-targeting qualities (*8*), which can be an advantage for clinical applications. AAVMYO2 and AAVMYO3 were less efficient than AAVMYO, but superior to AAV9, at transducing and therefore eliciting gene editing events in skeletal muscles (Fig. S6) as well as heart muscle (Fig. S7A). Notably, both AAVMYO2 and AAVMYO3, showed strong liver de-targeting (Fig. S7B-D). Nonetheless, because of the higher muscle tropism of AAVMYO, we opted to use this variant in further studies.

### AAVMYO-CRISPR/Cas9-mediated knockdown recapitulates conditional knockout model phenotypes for MuSK and myostatin/activin signaling

After successfully demonstrating the effectiveness of our model to perturb gene expression within skeletal muscle fibers, we asked whether we could recapitulate both loss and gain of muscle function phenotypes by targeting genes known to play a fundamental role in the regulation of muscle structure and growth.

We first chose to target the receptor tyrosine kinase MuSK, the signaling component of the Lrp4/MuSK receptor complex for motor neuron-released agrin (*21*). MuSK is essential for the formation and maintenance of the NMJ (*4, 22, 23*) and auto-antibodies against MuSK can cause myasthenia gravis (*24*), a disease leading to NMJ loss. As AAVMYO transduces all skeletal muscle fibers with high efficiency, we omitted tdTomato and instead focused on maximizing *Musk* gene editing, as no functional sgRNAs have been described. To this end, we inserted seven different sgRNAs into the constructs directed against exons localized in the 5’ region of the *Musk* gene. In a first set of experiments, we injected AAVMYO-7sgMusk (1.5 x 10^13^ vg/kg) or PBS (as a control) into the lateral tail vein of Cas9mKI (scheme Fig. 5A). By following the body weight, we noted that AAVMYO-7sgMusk-injected Cas9mKI started to lose weight after 14 days, reaching more than 20% at 20 days post injection (Fig. 5B). Their all- and forelimb grip strength was significantly lower than in controls (Fig. 5C, D) and they developed a severe kyphosis indicative of muscle weakness (Fig. 5E). Mice also showed signs of muscle fibrillation and ataxia, suggestive of denervation. To confirm this hypothesis, we measured mass in bulbar, fore- and hindlimb muscles. Indeed, all muscles of AAVMYO-7sgMusk-injected Cas9mKI were severely atrophic compared to controls (Fig 5F). CRISPR/Cas9 editing in TA muscle resulted in an almost complete loss of *Musk* mRNA expression (Fig. 5G). Whole-mount staining of the NMJ in the EDL muscle confirmed the loss of MuSK, which resulted in the very strong reduction of acetylcholine receptor (AChR) clusters (Fig. 5H). The presynaptic motor nerve terminals, visualized by a mixture of the SV2, directed against synaptic vesicle glycoprotein 2A, and 2H3, directed against the neurofilament-M protein, were still innervating the muscle fibers as in the controls (Fig. 5H). This loss of postsynaptic structures upon MuSK depletion is consistent with the results of transgenic mice deficient for Musk (*4, 22, 23*). Thus, our data show that the AAV-CRISPR/Cas9 system generates a somatic gene knockout whose phenotype is identical to germline-based methods.

**Figure 5:**
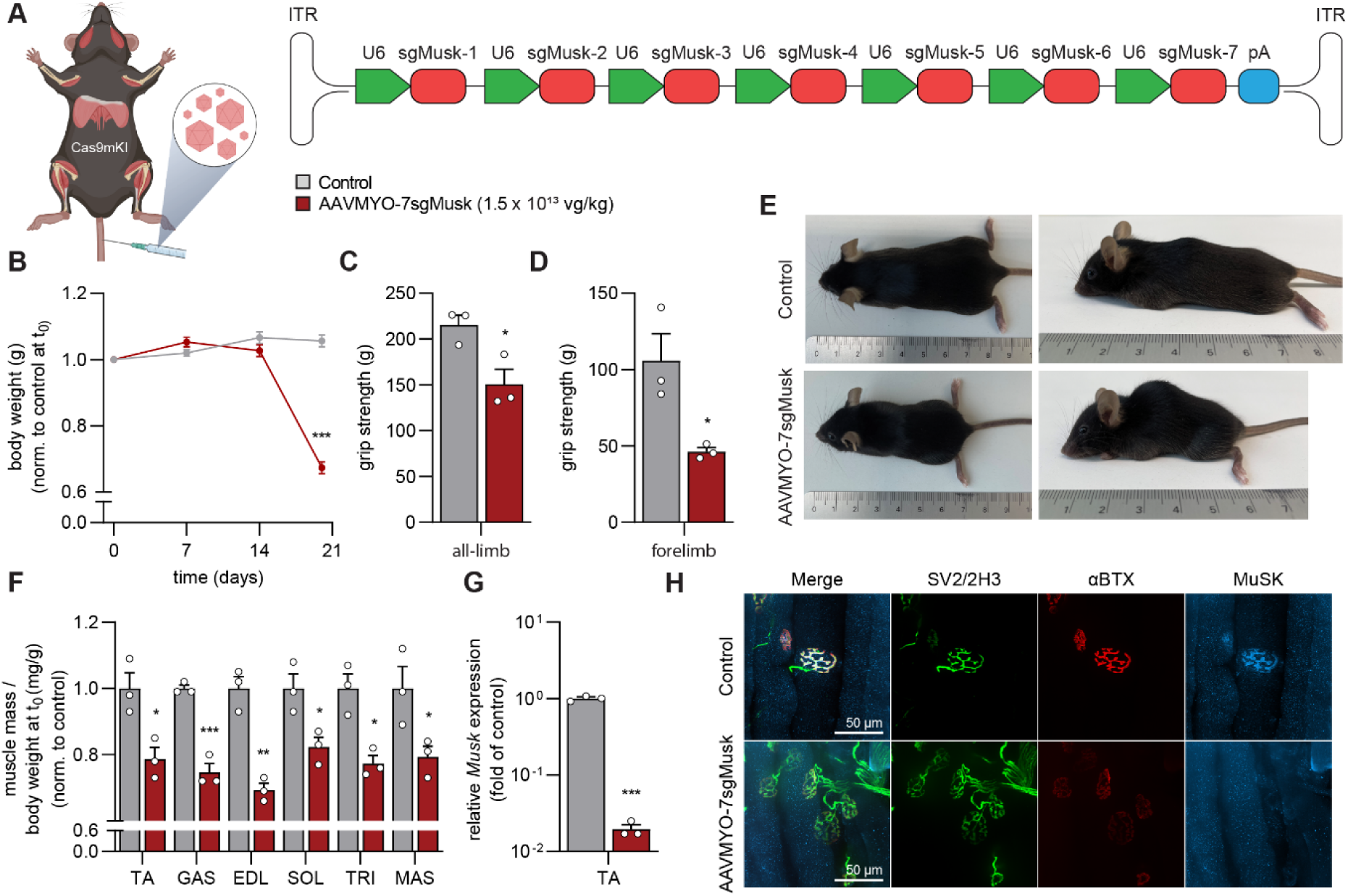
AAVMYO-CRISPR/Cas9 mediates systemic knockout of *Musk* and results in the loss of NMJs. (A) Schematic presentation of the experimental procedure. (B) Body weight progression of controls (grey) and AAVMYO-7sgMusk-injected Cas9mKI mice (red). All-limb (C) and forelimb (D) grip strength of control and AAVMYO-7sgMusk-injected Cas9mKI mice, 20 days post injection. (E) Representative photograph of control and AAVMYO-7sgMusk-injected Cas9mKI mice, 20 days post-injection. (F) Changes in mass of TA, GAS, EDL, SOL, TRI and *masseter* (MAS) muscle of AAVMYO-7sgMusk-injected Cas9mKI mice, compared to controls. (G) Relative mRNA expression of *Musk* in AAVMYO-7sgMusk- injected TA muscle of Cas9mKI mice. (H) Representative images of whole-mount preparations of EDL muscles of controls and Cas9mKI mice injected with AAVMYO-7sgMusk. The presynaptic nerve terminals are stained with a mixture of antibodies directed against synaptic vesicle glycoprotein 2A (SV2; yellow) and neurofilament (2H3; green). Fluorescently-labeled α-bungarotoxin (αBTX; red) was used to visualize postsynaptic AChRs. MuSK protein was stained using a specific antibody (blue). Data are means ± SEM. N = 3 mice. Statistical significance is based on unpaired t-test comparing to control. *P < 0.05, **P < 0.01, ***P < 0.001.

Next, we tested whether this system would also allow to restrict the depletion of MuSK to one or a few muscles. This might be advantage as loss of genes that are essential for muscle function (such as MuSK) will result in respiratory failure. As AAVMYO transduces the diaphragm well (see Fig. 4), respiratory failure may jeopardize the in depth analysis of limb muscles. To do this, we injected different doses of AAVMYO-7sgMusk into the right TA muscle of adult Cas9mKI mice (scheme Fig. 6A) and monitored the mice for 5 weeks. As controls, we chose to inject either AAVMYO-7sgMusk into wild-type (i.e. not expressing Cas9 in muscle) or injected PBS into Cas9mKI mice. As the two control conditions did not differ, we pooled data for further analysis (Fig. 6B-H). Despite intramuscular administration, mice receiving the highest dose of 3 x 10^11^ vg lost body mass (Fig. 6B) already 14 days after injection, reaching 20% body mass loss at 21 days and therefore requiring a humane endpoint. The second highest dose (1 x 10^11^ vg) induced measurable body mass losses by 28 days and reached our euthanization threshold at 35 days, while the two lowest doses did not induce body mass loss (Fig. 6B). Analysis of hindlimb muscle mass in the injected leg showed a dose-dependent decline in mass, which became significant compared to controls starting at a dose of 3.3 x 10^10^ vg (Fig. 6C). At the two highest doses (3 x 10^11^ and 1 x 10^11^ vg), significant loss of muscle mass was also observed in all contralateral, non-injected leg muscles compared to control mice (Fig. S8A). This highlights the high efficiency of the system, as the low amount of AAVMYO circulating in the blood upon intramuscular injection is sufficient to perturb gene function in remotely-positioned skeletal muscles (see also Fig. S4). At the two highest doses, *Musk* expression was significantly reduced down to 3.1% and 6.3% of control levels, respectively (Fig. 6D). *Musk* expression was also lower than in controls at a dose of 3.3 x 10^10^ and 1.1 x 10^10^ vg but both did not reach significance (Fig. 6D). As suggested by the overall loss of body and muscle mass, *Musk* mRNA abundance was significantly lower in the diaphragm (Fig. S8B) and the contralateral, non-injected TA muscle (Fig. S8C) compared to controls at the highest doses (8.9% of control in the diaphragm; 10.7% of control in TA) but not at the two lower doses. These results indicate that the lower dose of AAVMYO-7sgMusk largely restricts target knockdown to the injected muscle. To test for functional consequences by the loss of Musk upon local AAVMYO-7sgMusk injection, NMJ structure was examined in EDL muscle, which is adjacent to the TA muscle and hence becomes sufficiently transduced by intramuscular TA injection. Consistent with the phenotype of systemic MuSK depletion (Fig. 5H), postsynaptic AChR clusters were largely lost in AAVMYO-7sgMusk- injected Cas9mKI mice compared to controls, irrespective of dose (Fig. 6E). As a consequence of the loss of the postsynaptic structure, muscles become denervated, which causes re-expression of several synaptic genes along the entire muscle fiber (*20, 25*). To quantify the extent of denervation, we measured expression of mRNA coding for AChRα (*Chrna1*), the embryonic AChRγ subunit (*Chrng*) as well as growth arrest and DNA damage-inducible 45a (*Gadd45a*). The abundance of all transcripts was more than 10-times higher in the AAVMYO-7sgMusk-injected TA muscle of Cas9mKI mice than in controls (Fig. 6F-H). Consistent with whole-mount NMJ staining, the fold-induction of denervation-marker genes was independent of AAVMYO-7sgMusk dose. In summary, these experiments demonstrate that the lowering the dose of the injected AAV allows to perturb gene function largely restricted to the injected muscle without compromising the phenotype.

**Figure 6:**
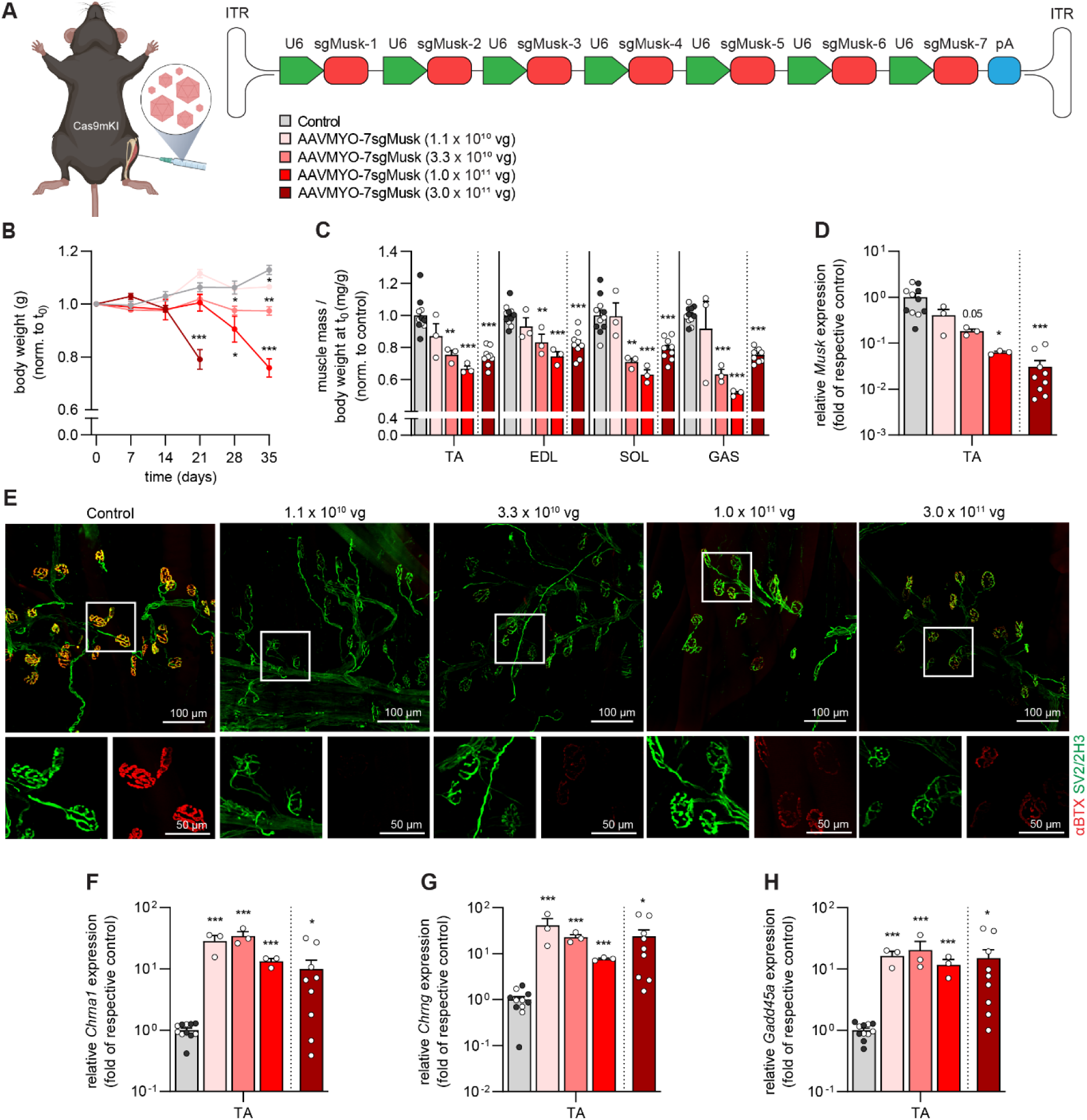
AAVMYO-CRISPR/Cas9 mediates local knockout of *Musk* and results in the loss of NMJs. (A) Schematic presentation of the experimental procedure using different amounts of sgRNA-delivering AAVMYO. (B) Body weight progression of controls (grey) and AAVMYO-7sgMusk-injected Cas9mKI mice (red colors) at the indicated doses. (C) Changes in muscle mass of AAVMYO-7sgMusk-injected Cas9mKI mice, compared to controls that included PBS-injected Cas9mKI mice (white dots) and AAVMYO-7sgMusk-injected wild-type mice (black dots). Values in the two control groups did not differ and were therefore combined. (D) Relative mRNA expression of *Musk* in AAVMYO-7sgMusk-injected TA muscle of Cas9mKI mice. (E) Representative whole-mount images of EDL muscles of controls and Cas9mKI mice injected with the indicated amount of AAVMYO-7sgMusk. The presynaptic nerve terminals are stained with a mixture of antibodies directed against synaptic vesicle glycoprotein 2A (SV2; yellow) and neurofilament (2H3; green). Fluorescently-labeled α-bungarotoxin (αBTX; red) was used to visualize postsynaptic AChRs. (F – H) Relative mRNA expression of denervation marker genes as indicated in AAVMYO-7sgMusk-injected TA muscles of Cas9mKI mice and controls. Note that Cas9mKI mice injected with the highest AAVMYO-7sgMusk dose were analyzed at 3 weeks post- injection while all the other mice were analyzed 5 weeks post-injection. Data are means ± SEM. N = 3- 11 mice. Statistical significance is based on unpaired t-test comparing to control. *P < 0.05, **P < 0.01, ***P < 0.001.

To test whether our system could also drive gain of muscle function, we next targeted myostatin (GDF-8), a TGF-β family protein secreted by skeletal muscle that acts as an inhibitor of muscle size (*26*). Deletion of *Mstn* in mice results in robust muscle hypertrophy (*27*) and naturally occurring *Mstn* null-mutants cause hypermuscularity in many species, including cows and humans (*28, 29*). Myostatin signals through a combination of type-2 and type-1 receptors. This signaling pathway is also activated by several other ligands, including activin. The two ligand-binding receptors are activin A receptor type- 2/IIA (ACVR2A or ACTRIIA) and type-2/IIB (AAVR2B or ACTRIIB). The activin A type-2 receptors are partially redundant as targeting both receptors elicits stronger muscle hypertrophy than deletion of each receptor individually (*30*). Upon ligand-binding, the type-2 receptors form a complex with type-1 activin A receptor-like kinase-4 (ALK4) and ALK5, which are also partially redundant, to trigger intracellular signaling.

To prevent partial compensation and to test the feasibility of the AAVMYO-CRISPR/Cas9 system to delete several genes, we targeted both mouse *Acvr2a* and *Acvr2b* genes by simultaneously injecting two AAVMYO viruses (each targeting one gene with seven different sgRNAs) at a dose of 3 x 10^11^ vg (each virus) into TA muscle of 8-week-old Cas9mKI mice (Fig. 7A) and analyzed muscles 6 weeks later. Virus-injected muscles expressed only 18% of *Acvr2a* and 26% or *Acvr2b* transcripts compared to PBS-injected muscle (Fig. 7B), confirming successful targeting. During the 6 weeks, AAVMYO-sgAcvr2a/b-injected mice gained significantly more body mass (Fig. 7C) and muscles of the injected leg were 40-50% heavier than in PBS-injected mice (Fig. 7D), highly comparable to the phenotype of *Acvr2a/b* double-knockout mice (*30*). Like in experiments targeting MuSK, the high dose intramuscular AAVMYO spread systemically, causing similar gains in muscle mass in the contralateral leg as the injected leg (Fig. S9A). Muscle growth in global *Mstn*-deficient mice is mediated *via* hyperplasia and hypertrophy (*27*), while myostatin signaling blockade after weaning (> 3-4 weeks) predominately stimulates hypertrophy (*26, 31, 32*). Consistent with these results, quantitative measurement of minimal fiber feret diameter (*33*) using immunohistochemistry in TA muscles injected with AAVMYO-sgAcvr2a/b (Fig. 7E) showed a consistent rightward shift in fiber size distribution and a significant increase in mean fiber size of all fiber types (Fig. 7F) without affecting fiber number (Fig. S9B). These results are highly consistent with those obtained with knockout mice and demonstrate the utility of AAVMYO-CRISPR/Cas9 to inactivate multiple genes and reproduce the phenotypes of traditional knockout mice, without the need to breed additional mouse lines.

**Figure 7:**
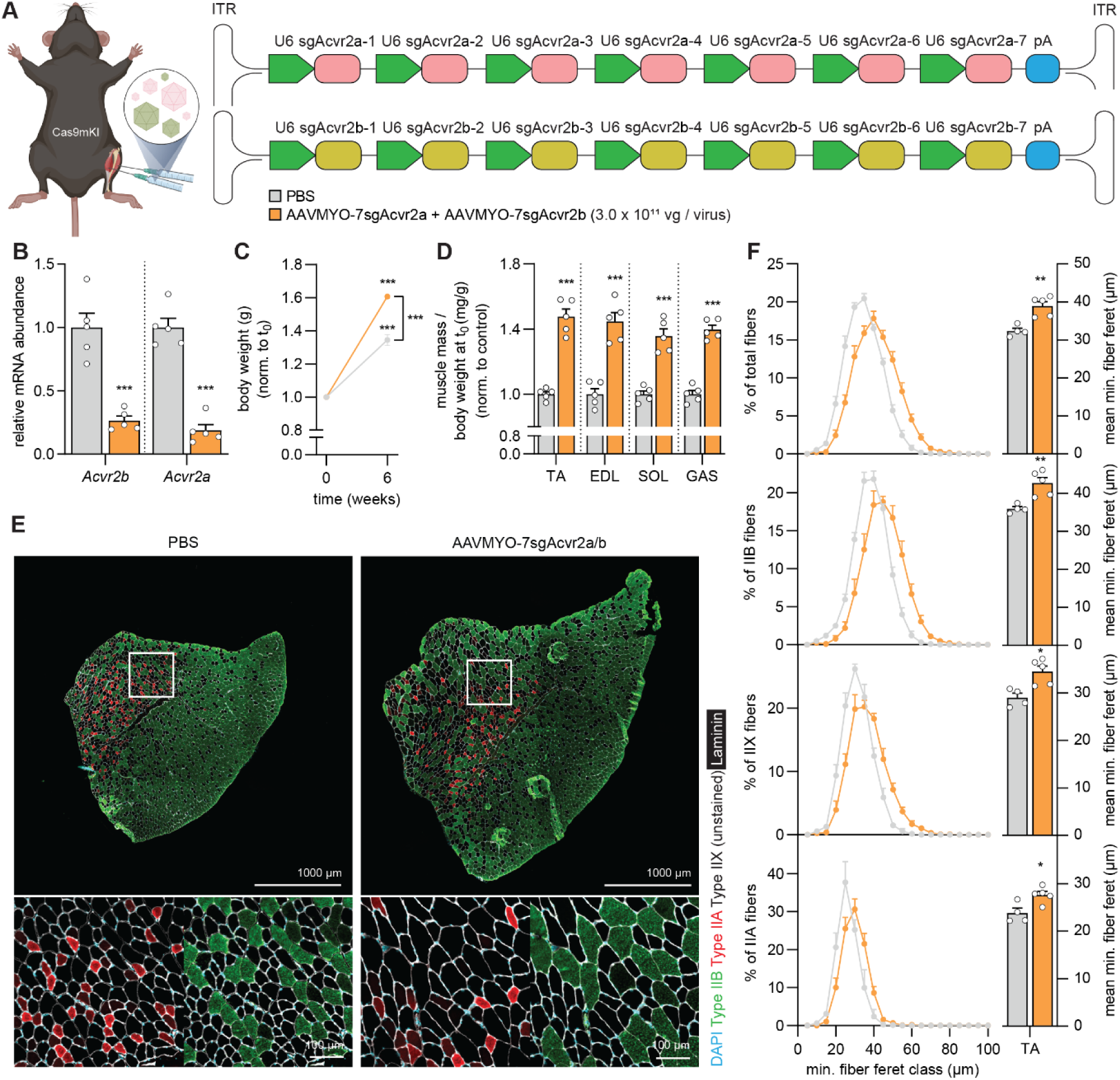
AAVMYO-CRISPR/Cas9-mediated double knockout of *Acvr2a*/*Acvr2b* causes strong skeletal muscle fiber hypertrophy. (A) Schematic representation of the experimental procedure. (B) Relative mRNA expression of *Acvr2a* and *Acvr2b* in *gastrocnemius* (GAS) muscle of Cas9mKI mice, injected with PBS or AAVMYO-7sgAcvr2a/b 6 weeks post-injection. (C) Body mass of Cas9mKI mice before and 6 weeks after intramuscular injection of PBS (grey) or AAVMYO-7sgAcvr2a/b (orange). (D) Mass of *tibialis anterior* (TA), *extensor digitorum longus* (EDL), *soleus* (SOL) and GAS muscles of the AAVMYO- 7sgAcvr2a/b-injected (orange) or PBS-injected (grey) legs of Cas9mKI mice. (E) Representative images of TA cross-sections stained with antibodies to type IIB (green), type IIA (red), laminin (white) and with DAPI (blue) from Cas9mKI mice injected as indicated. Note that type2X muscle fibers are not stained. (F) Total and fiber type-specific minimal fiber feret distribution (left) and mean minimal fiber feret (right) of TA muscle of Cas9mKI mice, injected with PBS (grey) or AAVMYO-7sgAcvr2a/b (orange). Data are means ± SEM. N = 4-5 mice. Statistical significance is based on unpaired t-test. *P < 0.05, **P < 0.01, ***P < 0.001.

## Discussion

This article presents a rapid and highly efficient tool to investigate the function of single or multiple genes in adult skeletal muscle fibers. Feasibility and efficiency of the system is demonstrated by knocking out essential genes for the integrity of the NMJ and skeletal muscle fiber growth.

We show that high, long-term Cas9 expression in skeletal muscle fibers does not affect muscle size or function. Others have used AAV to deliver Cre to LSL-Cas9KI mice to excise the stop cassette and drive Cas9 expression (*11, 34*). While this approach allows for the use of Cas9-GFP as a transfection marker and reduces any potential side effects of prolonged Cas9 expression, AAV-mediated delivery of Cre would also lead to Cas9 expression in any AAV-targeted tissues, including the heart and liver. To our best knowledge, highly specific AAV-compatible promoters for skeletal muscle fibers do not currently exist. As such, our AAVMYO-CRISPR/Cas9mKI strategy represents a major advancement for somatic gene perturbation of mouse skeletal muscle fibers.

While CRISPR/Cas9 systems for somatic gene deletion have been described for some tissues, including brain and liver (*11, 35*), such a versatile tool has so far been missing for skeletal muscle fibers. Previous work has demonstrated successful somatic gene editing using CRISPR in muscle stem cells (MuSCs) although with rather low efficiency (*36, 37*). While editing efficiency can be increased by sorting MuSCs based on a fluorescent transfection marker, this is not possible for multi-nucleated skeletal muscle fibers. Thus, the successful depletion of a gene by CRISPR in muscle fibers is only possible when indels are generated in both alleles in the majority of myonuclei. Such high efficiencies are not required in CRISPR/Cas9-mediated editing approaches that aim to correct gene mutations causing muscular dystrophies (*36, 38–42*). In these experiments, correcting the mutation in a subset of myonuclei and in one allele is sufficient as the corrected protein will distribute in a large part of the muscle fiber cytoplasm.

While CRISPR/Cas9-mediated gene deletion in mouse embryos has shortened the time to create founder mice to a few weeks and created the possibility to target multiple genes simultaneously (*43*), it still requires many founder breedings with different mouse lines to eventually achieve the final genotype needed for a study. The method we established here allows to conditionally knock out a single or multiple genes in muscle fibers without any prior breeding. We hypothesize that the use of mice expressing Cas9 at high levels in all muscle fibers in combination with AAVMYO to deliver multiple sgRNAs is key to achieve the efficient somatic gene deletions that mimic the phenotype of the respective knockout mouse. Our method also allows for both, systemic or local (in a single muscle) gene editing, which may be essential in cases where a gene knockout causes severe morbidity or death. Thus, this method also contributes to the 3R principle by strongly reducing the number of mice needed to investigate the function of genes *in vivo*.

The new method was established by targeting PKCα, as work in the retina has provided evidence for an efficient knockout using CRISPR/Cas9 (*15*) and based on the availability of high-quality antibodies to PKCα. Using this target, we were able to optimize the delivery method for the sgRNA by using AAVMYO instead of AAV9. With this optimized set-up, the amount of PKCα was lowered by approximately 80% in the injected muscle. Interestingly, DNA editing of the *Prkca* locus, measured by TIDE analysis, did not reach 80% but was only 23%. Several reasons can account for the quantitative difference between genome editing and loss of protein. First and foremost, only approximately 50% of the nuclei in a muscle isolate are myonuclei (*44, 45*). The remaining nuclei are derived from mono-nucleated cells, such as FAPs, macrophages, MuSCs, endothelial cells, smooth muscle cells or Schwann cells. All these non-muscle fiber cells do not express Cas9 in the Cas9mKi mice and are hence not edited. Hence, editing in muscle fibers would reach close to 50%. CRISPR/Cas9 editing in cultured C2C12 myotubes using the same sgRNA resulted in 60% editing and a loss of the protein of more than 90%. With this in mind (60% editing measured by TIDE analysis results in the almost complete loss of PKCα protein) and the fact that only 50% of the DNA in a muscle lysate are derived from myonuclei, the real *in vivo* editing efficiency would likely be above 75%. Another possible contributor to the incomplete editing efficiency may relate to the chromatin environment of the target site in the *Prkca* locus (*46*), which may differ between cultured C2C12 cells and muscle fibers *in vivo*. Since PKCα protein is mainly synthesized by muscle fibers (*10, 47*), the loss of the protein might be a better indicator for the efficiency of gene deletion.

As a functional proof-of-concept, we also perturbed MuSK function, which is essential for NMJ formation and maintenance (*21*). *Musk* expression in adult mice is confined to sub-synaptic nuclei, which lay directly underneath the NMJ. Sub-synaptic *Musk* expression is based on local, NMJ-derived signals that overwrite activity-mediated transcription suppression in non-synaptic myonuclei (*21*). Denervation and hence loss of electrical activity results in *Musk* re-expression in non-synaptic myonuclei. Hence, unlike *Prkca*, which is not specific to muscle fibers, efficient editing of myonuclear DNA should be sufficient to abrogate *Musk* expression in whole-muscle lysates. Indeed, *Musk* transcripts were reduced by 98% upon sgRNA expression. This strong reduction of *Musk* transcripts is likely due to the use of multiple sgRNAs that edit the *Musk* gene at multiple sites, which, in turn, may introduce large deletions that de-stabilize mRNA. Moreover, indels will result in frameshifts and the occurrence of premature termination codons that cause nonsense-mediated mRNA decay. A strong reduction of transcript levels was also observed for *Acvr2a* and *Acvr2b* using multiple sgRNAs.

Although the focus of our work was to use the MuSK knockdown as a proof-of-principle to demonstrate efficiency of the method, our data also show that MuSK is essential for the maintenance of the NMJ in the adult and that it is critical for muscle mass maintenance. This has so far only been shown indirectly by (i) injection of the MuSK ectodomain into adult mice that triggered the production of autoimmune antibodies and resulted in the deterioration of the NMJ reminiscent of myasthenia gravis (*48*), (ii) local shRNA-mediated suppression of *Musk* by electroporation, which led to NMJ loss (*4*) and (iii) by conditionally deleting *Musk* in muscle fibers by muscle creatine kinase-driven Cre, which caused death of the mice at approximately one month of age (*23*).

AAVMYO-CRISPR/Cas9 knockdown of *Musk*, *Acvr2a* and *Acvr2b* when injected systemically or at the highest dose into TA muscle resulted in a systemic loss of the targeted proteins. The systemic effect of the high intramuscular doses is likely based on the body-wide spreading of the sgRNA-expressing recombinant viruses *via* the blood stream and the subsequent transduction of skeletal muscles. Systemically administered AAVMYO targets all muscles but has the highest transduction rate in the diaphragm (Fig. 4C,(*7, 8*)). While such body-wide spreading may not be a problem for most experiments, in case of *Musk* deletion, NMJs deteriorate and muscles become denervated (*23*). At the highest dose of 3 x 10^11^ vg/mouse (corresponding to approx. 1.3 x 10^13^ vg/kg), mice started to lose weight 14 days post-injection and reached euthanization criteria (20% weight loss) at 3 weeks (Fig. 6B). Examination of the diaphragm muscle indicated NMJ deterioration. Based on this, mice injected with the highest dose needed to be analyzed already at 3 weeks post-injection, which explains the less severe phenotype in the hindlimbs. Lowering the dose of the injected virus to 3 or 1 x 10^10^ vg/mouse largely prevented weight and muscle mass loss in the contralateral leg while the injected muscle still showed all signs of NMJ deterioration and denervation. Thus, with the proper administration and viral titer, the AAVMYO-CRISPR/Cas9 method also allows for locally restricted perturbation of muscle function.

We also demonstrate efficacious, simultaneous inactivation of multiple genes (*Acvr2a* and *Acvr2b*) with this system, opening the possibility of studying several genes or signaling pathways concurrently. Although our experiments targeting *Prkca* indicate that one sgRNA can be sufficient to eliminate a gene, testing each sgRNA *in vitro* prior to *in vivo* application is laborious. Hence, we suggest targeting each gene with two to three different sgRNAs, minimizing the risk of insufficient protein loss. Since one AAV has sufficient packaging capacity for at least 7 sgRNAs, three genes can be silenced with one AAV. By delivering two AAVs (as done here for *Acvr2a* and *Acvr2b*), up to six independent genes could be silenced simultaneously, allowing interrogation of entire signaling pathways, specifically in skeletal muscle fibers.

In summary, we conclusively demonstrate that AAVMYO-mediated delivery of sgRNA to Cas9- expressing skeletal muscle fibers allows fast, efficient and specific gene knockouts. The multiplexable nature and capacity to induce systemic or local gene editing further strengthens the universality of the system. Therefore, this system provides an invaluable resource to perform loss-of-function studies in skeletal muscle fibers compared to traditional knockout mouse models and promises to greatly accelerate the interrogation of novel gene targets with a much reduced number of animals needed and thus will strongly contribute to our understanding of skeletal muscle biology.

## Material and Methods

### Mice

All procedures involving animals were performed in accordance with Swiss regulations and approved by the veterinary commission of the canton Basel Stadt. CRISPR/Cas9 knockin mice (*11*) were crossed with HSA-Cre (*13*) or HSA-Mer-Cre-Mer mice (*12*) to generate Cas9mKI or iCas9mKI, respectively. Littermates, knockin for Cas9 but not expressing Cre recombinase, were used as controls.

### Cell culture C2C12

Murine C2C12 myoblasts were cultured in growth medium (DMEM (Gibco) supplemented with 10% fetal bovine serum (Biological Industries) and 1% penicillin/streptomycin (Sigma)) at 37°C in an atmosphere of 5% CO_2_. After reaching 70% confluence, cells were transiently transfected using Lipofectamine 2000 (Invitrogen), according to the manufacturer’s protocol. At 48 h post-transfection, cells were incubated in growth medium, supplemented with 3 µg/ml puromycin (Sigma), for another 48 h to select for transfected cells. After selected cells reached confluence, cells were incubated in differentiation medium (DMEM (Gibco) supplemented with 2% horse serum (Biological Industries) and 1% penicillin/streptomycin for 5 days to induce formation of multinucleated myotubes.

### AAV administration

Prior to AAV administration, mice were anaesthetized by isoflurane inhalation. For intramuscular injection, the TA or TA and GAS muscle of adult mice (older than 6 weeks) was injected with 50 µL of AAV (3 x 10^11^ vg, if not stated differently) in PBS. For intravenous injection, 100 µL of AAV (1 x 10^14^ vg/kg) in PBS were injected into the lateral tail vein of 6-week-old mice. For targeting of PKCα or Acvr2a/Acvr2b, PBS or non-targeting AAV-injected control or Cas9mKI mice were used as control. For targeting of Musk, PBS-injected Cas9mKI mice or AAVMYO-7sgMusk-injected control mice were used as control.

### Denervation

Mice were anaesthetized by isoflurane inhalation 6 weeks post-AAV administration. After making a small incision on the skin between sciatic notch and knee, the sciatic nerve was exposed by gentle separation of muscles under the skin. The nerve was then lifted using a glass hook and disrupted by removing a 5 mm piece. The wound was closed by surgical clips and mice were returned to their cage. Mice were treated with Buprenorphine (0.1 mg/kg of body weight) one hour before and for two days after operation.

### sgRNA design and AAV vectors

The sgRNAs, listed in Table 1 were selected using CRISPOR (*19*) to minimize off-target effects and assembled as previously described using the multiplex CRISPR/Cas9 assembly kit (*49*). An array of three or seven human U6/sgRNA cassettes were cloned into an AAV transfer vectors. The AAV transfer vectors used for 3-plex sgRNA delivery into skeletal muscle were cloned between AAV serotype 2 ITR’s including a cloning site for multiplexed hU6-sgRNA insertions (MluI and KpnI (NEB)), the ubiquitous CMV promoter, tdTomato, WPRE and bovine growth hormone polyA signal. For 7-plex sgRNA delivery by AAV, the CMV-tdTomato-WPRE sequence was removed from the AAV transfer vector.

For in vitro CRISPR applications, sgRNAs were cloned into an all-in-one CRISPR/Cas9 vector using BbsI (NEB). The all-in-one CRISPR/Cas9 vector was cloned between AAV serotype 2 ITRs including a human U6 promoter, sgRNA scaffold, an EFS promoter, SpCas9 linked to puromycin N-acetyltransferase via a GSG-P2A linker and bovine growth hormone polyA signal. Complete vector maps and sequences are available upon request.

### AAV production, purification and titration

The AAV-sgRNA plasmid vectors were used for AAV production and purification. Briefly, adherent HEK293T cells were transiently transfected with transfer (AAV-sgRNA construct), AAV helper (AAV9 (a gift of J. M. Wilson Addgene, plasmid # 112867), AAVMYO (*7*), AAVMYO2 (*8*) or AAVMYO3 (*8*)) and pAdDeltaF6 helper (a gift from J. M. Wilson Addgene, plasmid # 112867) plasmid using PEI MAX (Polyscience). For small or large AAV preparations, ten or twenty HEK293T confluent 15 cm tissue culture plates were processed, respectively. The supernatant was collected 48 and 72 h post-transfection and cells were dislodged 72 hours post-transfection in PBS. Cells were centrifuged at 500g at 4°C for 10 min and resuspended in AAV lysis solution (50 mM Tris-HCl, 1 M NaCl, 10 mM NgCl_2_, pH 8.5). 50 U of salt active nuclease (Sigma) was added per harvested 15 cm dish and incubated at 37°C for 1 h with continuous shaking. The lysate was spun at 4000g at 4°C for 15 min and supernatant was collected. AAV particles from the supernatant were precipitated by adding polyethylene glycol 8000 (Sigma) to a final concentration of 8% (w/v), incubated for 2h at 4°C and then spun at 4000g at 4°C for 30 min. The supernatant was discarded, while the pellet was resuspended in AAV lysis buffer and pooled with the cell lysate. AAV particles were purified by using a 15-25-40-60% iodixanol (Serumwerk) gradient. The gradient was centrifuged at 63000 rpm (Beckman type 70 Ti rotor) for 2 h at 4°C and the AAV particles were collected from the 40-60% phase interface. The extract was passed through a 100 kDa MWCO filter (Millipore) and washed with PBS supplemented with 0.01% Pluronic F-68 surfactant (Gibco) until buffer was exchanged completely. The final volume was decreased to reach a final AAV concentration of > 1 x 10^13^ vg/ml. Virus was tittered using RT-qPCR targeted to the ITRs, as previously described (*50*), using a PvuII (NEB) -linearized plasmid standard. Primers used for titration are listed in Table 1.

### Protein isolation and Western blot analysis

Dissected muscles were snap-frozen in liquid nitrogen and pulverized. Proteins were extracted using RIPA lysis buffer (50 mM Tris-HCl pH 8.0, 150 mM NaCl, 1% NP-40, 0.5% sodium deoxycholate, 0.1% SDS) supplemented with protease and phosphatase inhibitors (both Roche) for 2 h at 4°C, followed by sonication. Lysates were centrifuged at 16000g for 20 min at 4°C and Pierce BCA Protein Assay Kit (Thermo Fisher Scientific) was used to determine cleared lysate concentration. Equalized protein samples were separated on 4-12% Bis-Tris Protein Gels (NuPage Novex), followed by transfer to nitrocellulose membranes (GE Healthcare Life Science). Membranes were blocked for 1 h by 5% BSA in PBS-T (1% Tween-20) and incubated with primary antibody in blocking solution overnight at 4°C. After 3 washes with PBS-T, membranes were incubated with secondary horseradish peroxidase-conjugated antibody for 1 h at RT. After washing 3 times with PBS-T, proteins were visualized using KPL LumiGLO (Seracare) and cheminumilescence was captured by a Fusion Fx machine (ViberLourmat). Protein abundance was quantified using the FusionCapt Advance software with linear background subtraction. Used antibodies are listed in Table 2.

**Table 2:** List of used antibodies

| Western Blot |  |  |  |
| --- | --- | --- | --- |
| PKCa | 1:1000 | 2056S | Cell Signaling |
| FLAG | 1:1000 | F3165 | Sigma |
| tdTomato | 1:5000 | orb182397 | Biorbyt |
| GFP | 1:1000 | 11814460001 | Roche |
| GAPDH | 1:5000 | 2118 | Cell Signaling |
| HDAC4 | 1:1000 | 7628S | Cell Signaling |
| Goat anti-rabbit HRP | 1:10000 | 111-035-003 | Jackson Immuno |
| Goat anti-mouse HRP | 1:10000 | 115-035-003 | Jackson Immuno |
| Donkey anti-goat HRP | 1:10000 | 705-035-003 | Jackson Immuno |
| Immunostaining |  |  |  |
| GFP | 1:400 | A10262 | Molecular Probes |
| MHC1 | 1:50 | BA-D5 | DSHB |
| MHC2a | 1:200 | SC-71 | DSHB |
| MHC2b | 1:100 | BF-F3 | DSHB |
| Laminin | 1:200 | L9393 | DSHB |
| Goat anti-mouse 405 | 1:50 | 115-475-207 | Jackson Immuno |
| Goat anti-mouse 568 | 1:200 | A-21124 | Invitrogen |
| Goat anti-mouse 488 | 1:200 | A-21042 | Invitrogen |
| Donkey anti-rabbit 647 | 1:300 | 711-605-152 | Jackson Immuno |
| Donkey anti-chicken 488 | 1:500 | 703-545-155 | Jackson Immuno |
| DAPI | 1:2000 | D9542 | Sigma |
| Whole mount immunostaining |  |  |  |
| Neurofilament | 1:2000 | 2H3 | DSHB |
| Synaptic vesicle protein | 1:400 | SV2 | DSHB |
| αBTX 647 | 1:500 | B-35450 | Invitrogen |
| Goat anti-mouse 555 | 1:500 | A-21127 | Invitrogen |

### Immunostaining of muscle cross sections and muscle histology analysis

Animal tissue was dissected, prepared and sectioned for immunohistochemistry as previously described (*51*). Tissue for analysis of cytosolic expression of GFP or tdTomato was directly fixed in ice-cold 4% PFA (Electron Microscopy Science) for 2 h at 4°, followed by dehydration with 20% sucrose (Sigma) in PBS at 4°C overnight. The next day, tissue was processed like non-fixed tissue as previously described (*51*). TA muscle cross sections were blocked and permeabilized for 30 min at RT with 3% BSA, 0.5% Triton X-100 in PBS. Primary antibodies were diluted in blocking solution for 2 h at RT. Sections were washed with PBS three times before being incubated in secondary antibody solution for 1 h at RT. All antibodies are listed in Table 2. Sections were washed with PBS four times and mounted with ProLong Gold Antifade Mountant (Invitrogen). Muscle sections were imaged at the Biozentrum Imaging Core Facility with a SpinD confocal microscope (Olympus). The previously described script for automated muscle cross section analysis (*52*) was further developed in-house and is available upon request.

### Whole-mount NMJ staining

EDL muscles were fixed, cut into bundles and prepared for NMJ staining as previously described (*51*). The presynapse was visualized using a primary antibody mix against neurofilament and synaptic vesical protein, while the postysnapse was stained using A647-conjugated α-bungarotoxin. NMJs were imaged at the Biozentrum Imaging Core Facility with a SpinD confocal microscope (Olympus).

### In vitro muscle force

Fast-twitch EDL and slow-twitch SOL muscles were carefully isolated for in vitro force and fatigue test as previously described (*51*). The measurement was carried out on the 1200 A Isolated Muscle System (Aurora Scientific) in Ringer solution (137 mM NaCl, 24 mM NaHCO_3_, 11 mM glucose, 5 mM KCl, 2 mM CaCl_2_, 1 mM MgSO_4_, 1 mM NaH_2_PO_4_) which was gassed with 95% O_2_, 5% CO_2_ and kept at 30 °C.

### Genomic DNA isolation, PCR amplification and TIDE

Cells were washed in PBS, while dissected tissue was snap-frozen in liquid nitrogen and pulverized in liquid nitrogen. Genomic DNA from cells or tissue was isolated using the DNeasy blood and Tissue kit (Qiagen) according to the manufacturer’s protocol. DNA was amplified using standard PCR using LongAmp Taq DNA polymerase (NEB) targeting the CRISPR locus with a 200-500 bp long amplicon. Used primers are listed in Table 1. PCR amplicons were purified using AMPure XP beads (Beckman) and Sanger-sequenced using one of the two PCR primers (Microsynth). TIDE was applied to sequencing chromatograms to assess CRISPR editing efficiency of the target locus (*53*).

### Amplicon deep sequencing analysis

PCR of genomic DNA was performed using LongAmp Taq DNA polymerase (NEB) and primers designed against an amplicon of 200-500 bp targeting the CRISPR-locus and the top four CRISPOR-predicted off-targets. First-round PCR primers contained adapter sequence for DNA/RNA UD Indexes (Illumina). The second round of PCR and pooling of samples was performed according to the Illumina Nextera DNA library prep guide. Pooled amplicons were sequenced with standard 500 cycles kit PE 2x251 on an Illumina MiSeq instrument. Samples were demultiplexed according to assigned barcodes and FASTQ files were analyzed using the CRISPResso2 software package (*54*).

### AAV genome copy number quantification

The AAV viral genome copy number per nuclei was determined by RT-qPCR using PowerUp SYBR Green Master Mix (Applied Biosystems) and primers (Table 1) targeting the tdTomato-WPRE sequence (AAV) or the R26 locus (nuclei) on a QuantStudio5 (Applied Biosystems) instrument. The cycle threshold (CT) values were converted into copy numbers by measuring against a standard curve of the AAV transfer plasmid or the Ai9 plasmid (a gift from H. Zeng Addgene plasmid # 22799). The AAV genome copy number was divided by the number of nuclei to normalize for tissue input.

### RT-qPCR

Pulverized muscle tissue was lysed in RLT buffer (Qiagen) and RNA was extracted using the RNeasy® Mini Kit for fibrous tissue (Qiagen). cDNA was reverse transcribed using the iScript^TM^ cDNA synthesis kit (Bio-Rad) and 500 ng of RNA according to the manual. RT-qPCR was performed using PowerUp SYBR Green Master Mix (Applied Biosystems) and target specific primers (table 1) on a QuantStudio5 (Applied Biosystems) instrument. Data were analyzed using the comparative CT method (2^−ΔΔCq^). Raw CT values of targets were normalized to CT values of a housekeeper (Tata-box-binding protein), which was stable between conditions, and then further normalized to the control group for visualization.

### Statistical analysis

All values are expressed as mean +/- SEM, unless stated otherwise. Data were analyzed in GraphPad Prism 8. Unpaired Student t tests were used for pairwise comparison. One-way ANOVAs with Fisher’s LSD post-hoc tests were used to compare between three groups, while Tukey post-hoc tests were used for comparison between more than three groups, so long as the ANOVA reached statistical significance. Significant differences (*P < 0.05, **P < 0.01, *** P < 0.001) are reported on figures, where appropriate.

## Acknowledgments

We thank the Biozentrum core facilities for their technical support with imaging (Imaging Core Facility), sequencing (Genomics Facility), computing (Scicore) and mice housing (Animal Core Facility). We thank Dr. D.J. Ham for his comments on the manuscript.

## Funding

This work was supported by funds from:

- Swiss National Science Foundation 189248
- The cantons of Basel-Stadt and Basel-Landschaft

## Author contributions

Conceptualization: MT, MAR; methodology: MT; investigations: MT, SL, FO; statistical analysis: MT; visualization: MT; supervision: MAR, RJP, DG; writing original draft: MT, MAR; writing review and editing: MT, DG, MAR; funding acquisition: MAR

Corresponding author: MAR

## Competing interests

Authors declare that they have no competing interests.

## Data and materials availability

All data are available in the main text or the supplementary materials.

## Notes

### Competing Interest Statement

The authors have declared no competing interest.

## References

1. M. Adli, The CRISPR tool kit for genome editing and beyond. Nature communications 9, 1911 (2018).

2. J. C. Bruusgaard, K. Liestol, M. Ekmark, K. Kollstad, K. Gundersen, Number and spatial distribution of nuclei in the muscle fibres of normal mice studied in vivo. J Physiol 551, 467–478 (2003).

3. B. Hu et al., Therapeutic siRNA: state of the art. Signal Transduct Target Ther 5, 101 (2020).

4. X. C. Kong, P. Barzaghi, M. A. Ruegg, Inhibition of synapse assembly in mammalian muscle in vivo by RNA interference. EMBO Rep 5, 183–188 (2004).

5. M. Sandri et al., Foxo transcription factors induce the atrophy-related ubiquitin ligase atrogin-1 and cause skeletal muscle atrophy. Cell 117, 399–412 (2004).

6. D. R. Bisset et al., Therapeutic impact of systemic AAV-mediated RNA interference in a mouse model of myotonic dystrophy. Hum Mol Genet 24, 4971–4983 (2015).

7. J. Weinmann et al., Identification of a myotropic AAV by massively parallel in vivo evaluation of barcoded capsid variants. Nature communications 11, 5432 (2020).

8. J. El Andari et al., Semirational bioengineering of AAV vectors with increased potency and specificity for systemic gene therapy of muscle disorders. Sci Adv 8, eabn4704 (2022).

9. M. Tabebordbar et al., Directed evolution of a family of AAV capsid variants enabling potent muscle-directed gene delivery across species. Cell 184, 4919–4938 e4922 (2021).

10. M. J. Petrany et al., Single-nucleus RNA-seq identifies transcriptional heterogeneity in multinucleated skeletal myofibers. Nature communications 11, 6374 (2020).

11. R. J. Platt et al., CRISPR-Cas9 knockin mice for genome editing and cancer modeling. Cell 159, 440–455 (2014).

12. J. J. McCarthy, R. Srikuea, T. J. Kirby, C. A. Peterson, K. A. Esser, Inducible Cre transgenic mouse strain for skeletal muscle-specific gene targeting. Skelet Muscle 2, 8 (2012).

13. M. Leu, et al., Erbb2 regulates neuromuscular synapse formation and is essential for muscle spindle development. Development (Cambridge, England) 130, 2291–2301 (2003).

14. V. Thomanetz et al., Ablation of the mTORC2 component rictor in brain or Purkinje cells affects size and neuron morphology. J Cell Biol 201, 293–308 (2013).

15. S. Sarin et al., Role for Wnt Signaling in Retinal Neuropil Development: Analysis via RNA-Seq and In Vivo Somatic CRISPR Mutagenesis. Neuron 98, 109–126 e108 (2018).

16. C. F. Bentzinger et al., Skeletal muscle-specific ablation of raptor, but not of rictor, causes metabolic changes and results in muscle dystrophy. Cell Metab 8, 411–424 (2008).

17. C. L. Bell et al., The AAV9 receptor and its modification to improve in vivo lung gene transfer in mice. J Clin Invest 121, 2427–2435 (2011).

18. H. Zhu, T. Wang, R. John Lye, B. A. French, B. H. Annex, Neuraminidase-mediated desialylation augments AAV9-mediated gene expression in skeletal muscle. J Gene Med 20, e3049 (2018).

19. J. P. Concordet, M. Haeussler, CRISPOR: intuitive guide selection for CRISPR/Cas9 genome editing experiments and screens. Nucleic Acids Res 46, W242–W245 (2018).

20. H. Lin et al., Decoding the transcriptome of denervated muscle at single-nucleus resolution. J Cachexia Sarcopenia Muscle 13, 2102–2117 (2022).

21. L. A. Tintignac, H. R. Brenner, M. A. Ruegg, Mechanisms Regulating Neuromuscular Junction Development and Function and Causes of Muscle Wasting. Physiol Rev 95, 809–852 (2015).

22. T. M. DeChiara et al., The receptor tyrosine kinase MuSK is required for neuromuscular junction formation in vivo. Cell 85, 501–512 (1996).

23. B. A. Hesser, O. Henschel, V. Witzemann, Synapse disassembly and formation of new synapses in postnatal muscle upon conditional inactivation of MuSK. Mol Cell Neurosci 31, 470–480 (2006).

24. W. Hoch et al., Auto-antibodies to the receptor tyrosine kinase MuSK in patients with myasthenia gravis without acetylcholine receptor antibodies. Nature medicine 7, 365–368 (2001).

25. V. Witzemann, H. R. Brenner, B. Sakmann, Neural factors regulate AChR subunit mRNAs at rat neuromuscular synapses. J Cell Biol 114, 125–141 (1991).

26. S. J. Lee, Myostatin: A Skeletal Muscle Chalone. Annu Rev Physiol, (2022).

27. A. C. McPherron, A. M. Lawler, S. J. Lee, Regulation of skeletal muscle mass in mice by a new TGF-beta superfamily member. Nature 387, 83–90 (1997).

28. L. Grobet et al., A deletion in the bovine myostatin gene causes the double-muscled phenotype in cattle. Nat Genet 17, 71–74 (1997).

29. M. Schuelke et al., Myostatin mutation associated with gross muscle hypertrophy in a child. N Engl J Med 350, 2682–2688 (2004).

30. S. J. Lee et al., Functional redundancy of type I and type II receptors in the regulation of skeletal muscle growth by myostatin and activin A. Proc Natl Acad Sci U S A 117, 30907–30917 (2020).

31. S. Welle, K. Bhatt, C. A. Pinkert, R. Tawil, C. A. Thornton, Muscle growth after postdevelopmental myostatin gene knockout. Am J Physiol Endocrinol Metab 292, E985–991 (2007).

32. H. Amthor et al., Muscle hypertrophy driven by myostatin blockade does not require stem/precursor-cell activity. Proc Natl Acad Sci U S A 106, 7479–7484 (2009).

33. A. Briguet, I. Courdier-Fruh, M. Foster, T. Meier, J. P. Magyar, Histological parameters for the quantitative assessment of muscular dystrophy in the mdx-mouse. Neuromuscul Disord 14, 675–682 (2004).

34. R. D. Chow et al., AAV-mediated direct in vivo CRISPR screen identifies functional suppressors in glioblastoma. Nature neuroscience 20, 1329–1341 (2017).

35. T. Katsuda et al., Rapid in vivo multiplexed editing (RIME) of the adult mouse liver. Hepatology, (2022).

36. M. Tabebordbar et al., In vivo gene editing in dystrophic mouse muscle and muscle stem cells. Science 351, 407–411 (2016).

37. L. He et al., CRISPR/Cas9/AAV9-mediated in vivo editing identifies MYC regulation of 3D genome in skeletal muscle stem cell. Stem Cell Reports 16, 2442–2458 (2021).

38. L. Amoasii et al., Single-cut genome editing restores dystrophin expression in a new mouse model of muscular dystrophy. Sci Transl Med 9, (2017).

39. D. U. Kemaladewi et al., Correction of a splicing defect in a mouse model of congenital muscular dystrophy type 1A using a homology-directed-repair-independent mechanism. Nature medicine 23, 984–989 (2017).

40. Y. Zhang et al., Enhanced CRISPR-Cas9 correction of Duchenne muscular dystrophy in mice by a self-complementary AAV delivery system. Sci Adv 6, eaay6812 (2020).

41. C. E. Nelson et al., In vivo genome editing improves muscle function in a mouse model of Duchenne muscular dystrophy. Science 351, 403–407 (2016).

42. L. Amoasii et al., Gene editing restores dystrophin expression in a canine model of Duchenne muscular dystrophy. Science 362, 86–91 (2018).

43. N. S. McCarty, A. E. Graham, L. Studena, R. Ledesma-Amaro, Multiplexed CRISPR technologies for gene editing and transcriptional regulation. Nature communications 11, 1281 (2020).

44. H. Schmalbruch, U. Hellhammer, The number of nuclei in adult rat muscles with special reference to satellite cells. Anat Rec 189, 169–175 (1977).

45. M. Bengtsen et al., Comparing the epigenetic landscape in myonuclei purified with a PCM1 antibody from a fast/glycolytic and a slow/oxidative muscle. PLoS Genet 17, e1009907 (2021).

46. A. M. Chakrabarti et al., Target-Specific Precision of CRISPR-Mediated Genome Editing. Mol Cell 73, 699–713 e696 (2019).

47. M. Dos Santos et al., Single-nucleus RNA-seq and FISH identify coordinated transcriptional activity in mammalian myofibers. Nature communications 11, 5102 (2020).

48. A. R. Punga, M. Maj, S. Lin, S. Meinen, M. A. Ruegg, MuSK levels differ between adult skeletal muscles and influence postsynaptic plasticity. Eur J Neurosci 33, 890–898 (2011).

49. T. Sakuma, A. Nishikawa, S. Kume, K. Chayama, T. Yamamoto, Multiplex genome engineering in human cells using all-in-one CRISPR/Cas9 vector system. Sci Rep 4, 5400 (2014).

50. F. Wang, X. Cui, M. Wang, W. Xiao, R. Xu, A reliable and feasible qPCR strategy for titrating AAV vectors. Medical science monitor basic research 19, 187–193 (2013).

51. D. J. Ham et al., The neuromuscular junction is a focal point of mTORC1 signaling in sarcopenia. Nature communications 11, 4510 (2020).

52. L. Encarnacion-Rivera, S. Foltz, H. C. Hartzell, H. Choo, Myosoft: An automated muscle histology analysis tool using machine learning algorithm utilizing FIJI/ImageJ software. PLoS One 15, e0229041 (2020).

53. E. K. Brinkman, T. Chen, M. Amendola, B. van Steensel, Easy quantitative assessment of genome editing by sequence trace decomposition. Nucleic Acids Res 42, e168 (2014).

54. K. Clement et al., CRISPResso2 provides accurate and rapid genome editing sequence analysis. Nat Biotechnol 37, 224–226 (2019).

