## Supplementary Material for "Fast, multiplexable and highly efficient somatic gene deletions in adult mouse skeletal muscle fibers using AAV-CRISPR/Cas9"

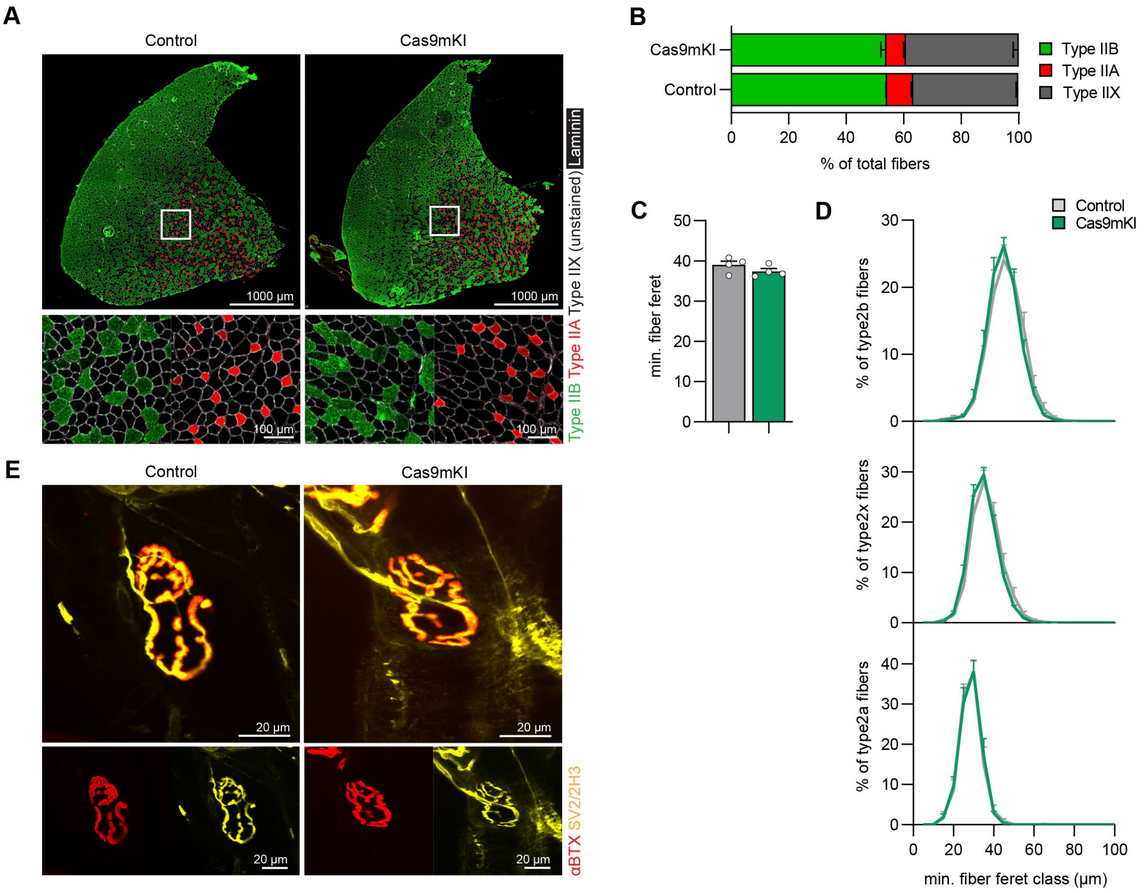


**Figure S1: Validation of Cas9mKI mice.** (A) Cross-sections of *tibialis anterior* (TA) muscle stained for type IIB (green), type IIA (red) fibers and laminin (white) from control and Cas9mKI mice. (B) Quantification of the TA fiber type composition. (C) Mean minimal fiber feret of TA muscle. (D) Minimal fiber feret distribution of TA muscle of control or Cas9mKI mice according to their fiber type. (E) Whole-mounts of the neuromuscular junctions of the *extensor digitorum longus* (EDL) muscle stained for the nerve terminal with a mixture of antibodies directed against synaptic vesicle glycoprotein 2A (SV2; yellow) and neurofilament (2H3; yellow) and for the postsynaptic AChRs with α-bungarotoxin (αBGT; red). Data are means ± SEM with n = 4 mice for each genotype. None of the data in B – D are significantly different between control and Cas9mKI mice ($P > 0.05$) using unpaired t-test.

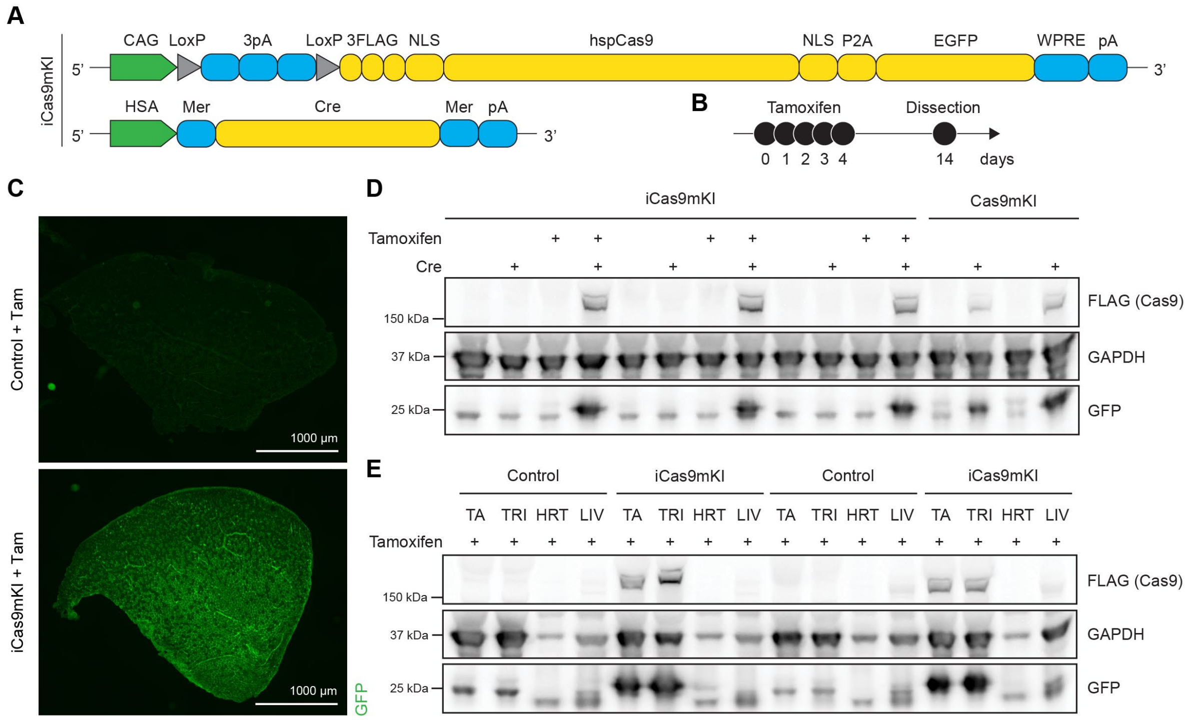


**Figure S2: Validation of iCas9mKI mice.** (A) Schematic of the iCas9mKI mouse model. For abbreviations, see legend to Figure 1. Mer: mutated estrogen receptor. (B) Timeline of tamoxifen injection and mouse analysis 14 days post-injection. (C) Cross-sections from *tibialis anterior* (TA) muscle from control and iCas9mKI mice stained for GFP. (D) Western blot analysis of TA muscle for the FLAG-tag and GFP in iCas9mKI in Cre-positive or Cre-negative mice upon tamoxifen injection. Glyceraldehyde-3-phosphate dehydrogenase (GAPDH) was used as loading control. (E) Western blot analysis for FLAG-tag and GFP in TA, *triceps brachii* (TRI), heart (HRT) and liver (LIV) after tamoxifen administration in control and iCas9mKI mice.

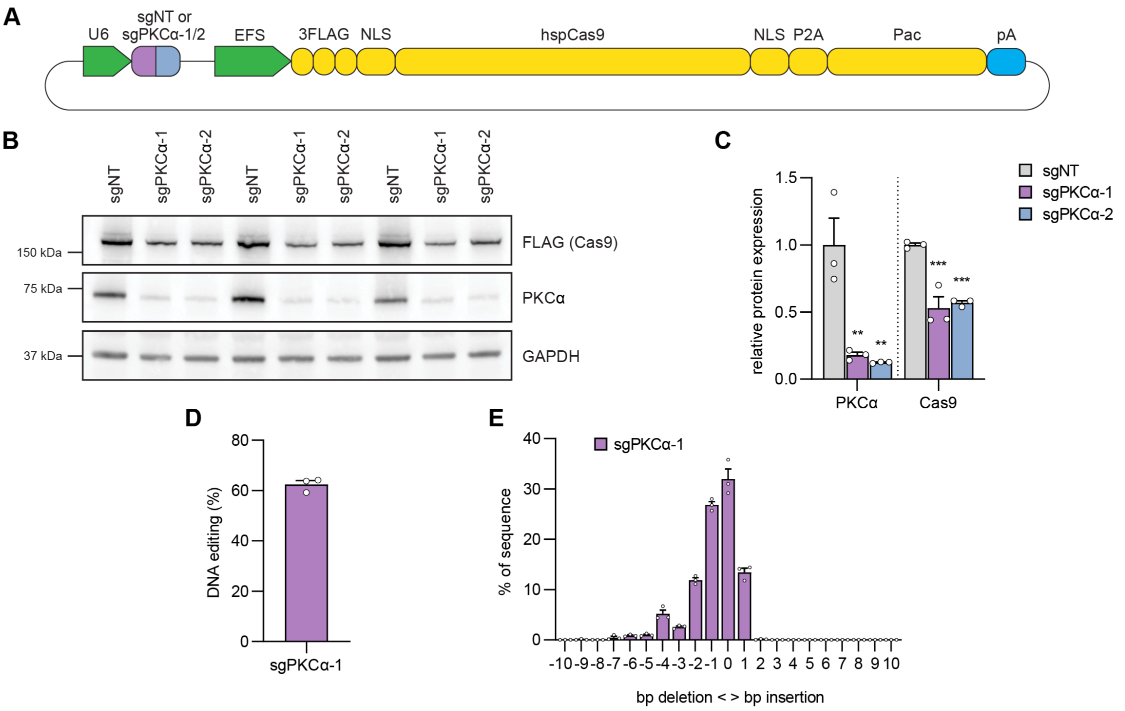


**Figure S3: Efficacy testing of sgRNAs in cultured C2C12 myotubes.** (A) Schematic of the expression plasmid used for transfection of C2C12 myoblasts. For abbreviations, see legend to Figure 1. U6: human U6 promoter; EFS: eukaryotic translation elongation factor 1 α short promoter; Pac: puromycin N-acetyltransferase. (B) Western blot analysis for FLAG-tagged Cas9 and PKCα using lysates from C2C12 myotubes after transfection with the indicated constructs and puromycin selection. (C) Quantification of protein abundance. (D) Tracking of Indels by Decomposition (TIDE) analysis of sgPKCα-1-expressing cells. (E) Frequency distribution of the DNA editing events around the binding site of sgPKCα-1. Data are means ± SEM with n = 3 wells for each sgRNA. Significance was determined using one-way ANOVA with Fishers LSD post-hoc test. *P < 0.05, **P< 0.01, ***P < 0.001.

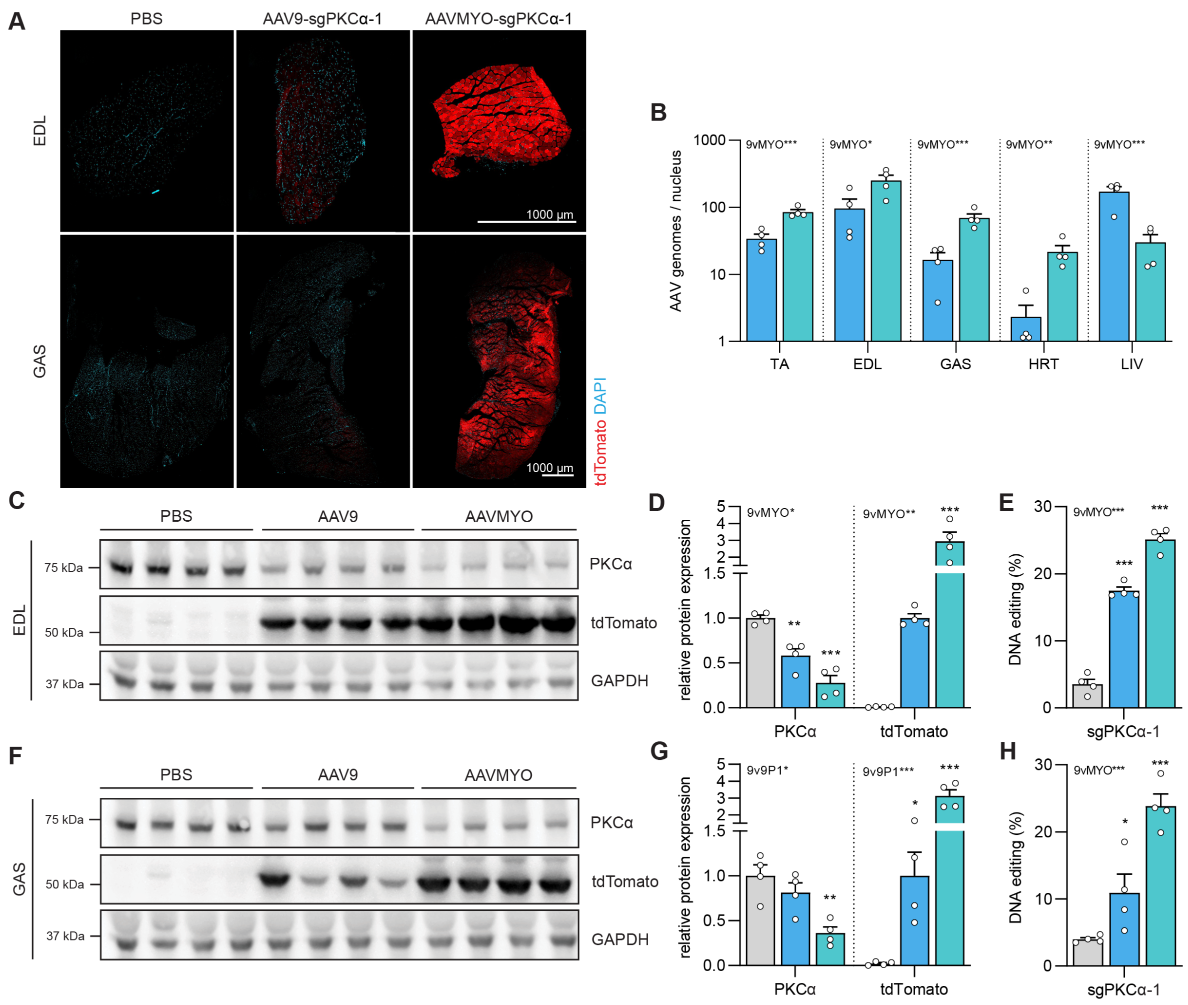


**Figure S4: AVVMYO-mediated sgRNA delivery into TA muscle induces a strong reduction of PKCα in nearby muscles.** (A) Representative images of cross-sections of *extensor digitorum longus* (EDL) and *gastrocnemius* (GAS) muscle stained for tdTomato (red) and DAPI (blue). (B) Distribution of AAVs in TA, EDL, GAS, heart (HRT) and liver (LIV) upon intramuscular injection of AAV9-sgPKCα-1 (light blue) or AAVMYO-sgPKCα-1 (cyan) into Cas9mKI mice. (C, F) Western blot analysis for PKCα and tdTomato of the indicated muscles and conditions. (D, G) Quantification of Western blots shown in C, F for the indicated proteins. Results for PKCα were normalized to the levels in PBS-injected muscles (grey). For tdTomato, levels of AAV9-sgPKCα-1-injected muscle were set to 1. (E, H) Total INDEL formation analysis by TIDE. Data are means ± SEM. N = 4 mice for each condition. Significance was determined using one-way ANOVA with Fishers LSD post-hoc test (D, E, G, H) or unpaired t-test (B). *P < 0.05, **P < 0.01, ***P < 0.001.

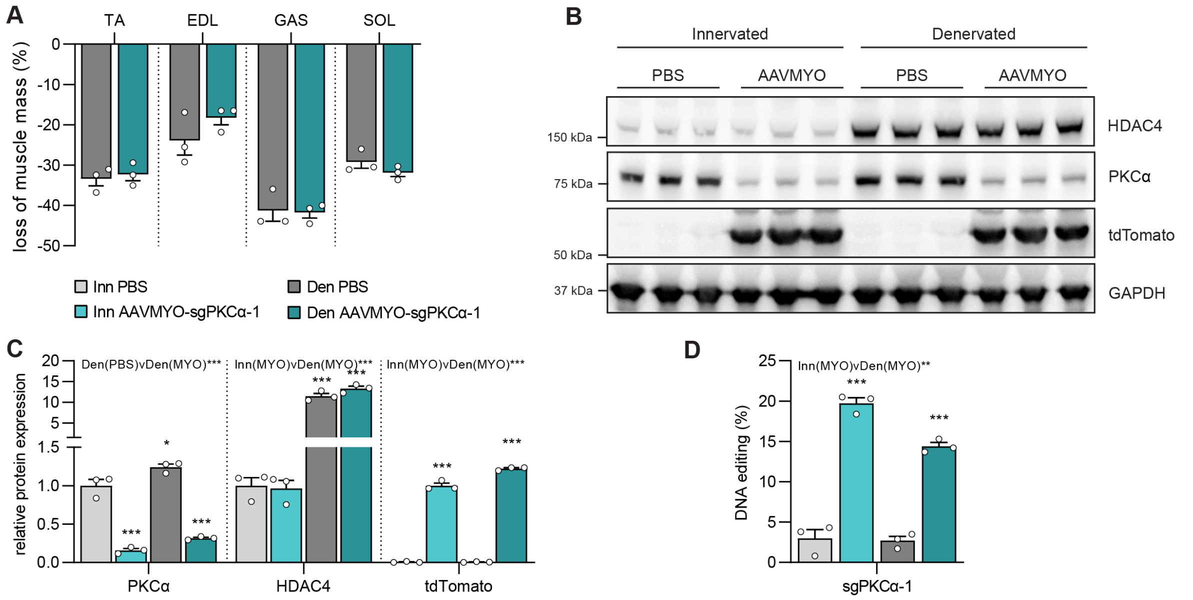


**Figure S5: AAVMYO-CRISPR/Cas9-induced loss of *Prkca* in denervated muscle.** (A) Loss of mass after 14 days of denervation of *tibialis anterior* (TA), *extensor digitorum longus* (EDL), *gastrocnemius* (GAS) and *soleus* (SOL) muscles after normalization to the innervated, contralateral muscle in each animal. (B) Western blot analysis and (C) quantification of protein abundance of PKCα, HDAC4 and tdTomato in either innervated or denervated TA muscle of Cas9mKI mice injected with PBS or AAVMYO-sgPKCα-1. (D) INDEL analysis by TIDE in the different conditions. Data are means ± SEM. N = 3 mice for each condition. Significance was determined using one-way ANOVA with Tukey's post-hoc test. *P < 0.05, **P < 0.01, ***P < 0.001.

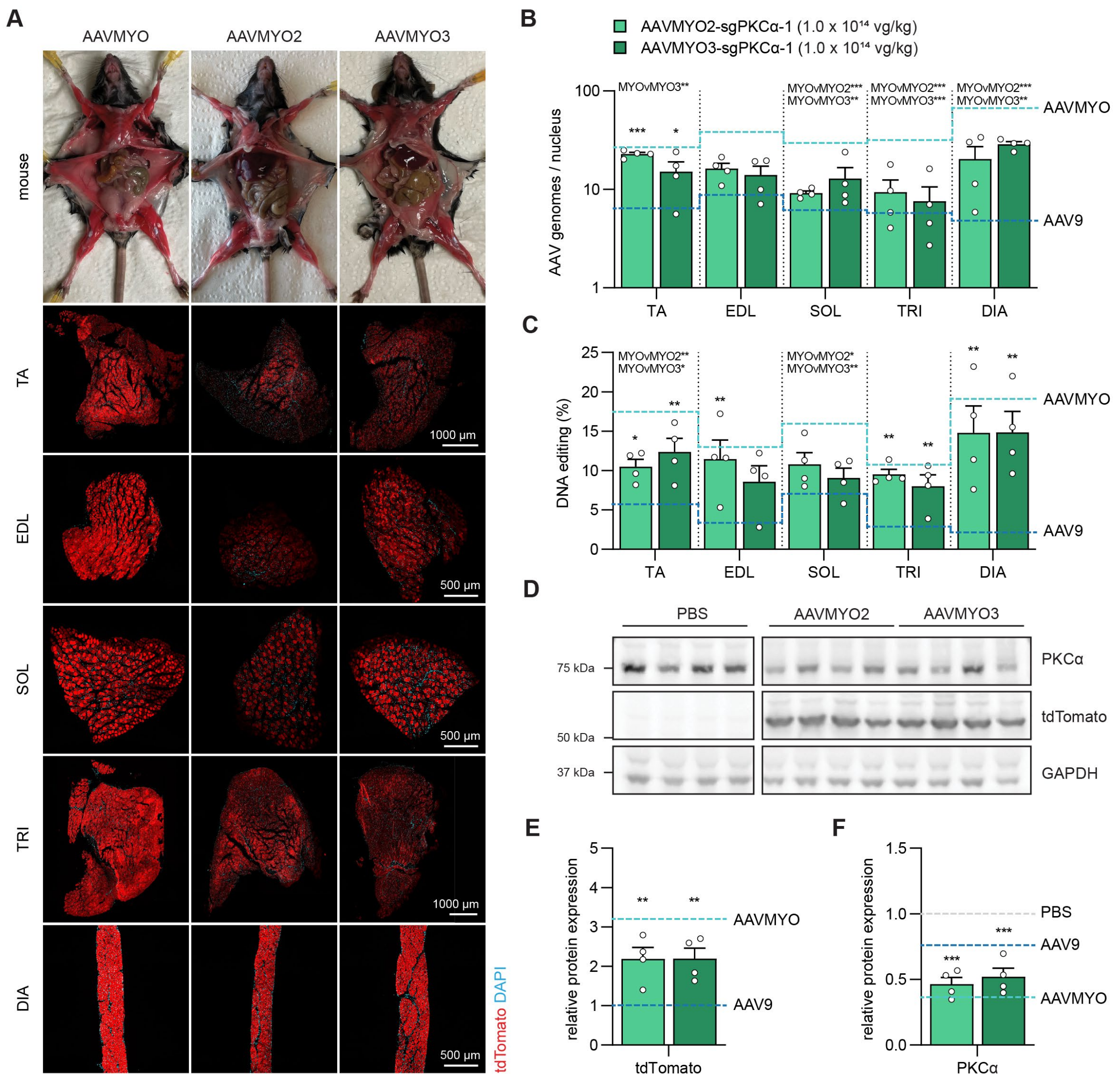


**Figure S6: Efficacy of sgRNA delivery using the liver-detargeted AAVMYO2 and AAVMYO3 compared to AAVMYO.** (A) Representative images of dissected mice and cross-sections of *tibialis anterior* (TA), *extensor digitorum longus* (EDL), *soleus* (SOL), *triceps brachii* (TRI) or diaphragm (DIA) muscle stained for tdTomato (red) and DAPI (blue), 6 weeks post-intravenous injection of PBS or AAV (1.0 x $10^{14}$ vg/kg) into Cas9mKI mice. For comparison, images of AAVMYO- sgPKCα-1-injected mice are included, same as on fig. 4B. (B) Distribution of AAVs in TA, EDL, SOL, TRI and DIA upon intravenous injection of AAVMYO2-sgPKCα-1 (light green) or AAVMYO3-sgPKCα-1 (dark green) into Cas9mKI mice. (C) Total INDEL formation analysis by TIDE. (D) Western blot analysis and its quantification (E, F) for PKCα (E) and tdTomato (F) in TA muscle of Cas9mKI mice injected with AAVMYO2 or AAVMYO3. Dashed lines in B, C, E and F show mean values for AAV9 (light blue), AAVMYO (cyan) and PBS (grey) when appropriate. Data are means ± SEM. N = 4 - 5 mice for each condition. Significance was determined using one-way ANOVA with Tukey's post-hoc test. *P < 0.05, **P < 0.01, ***P < 0.001.

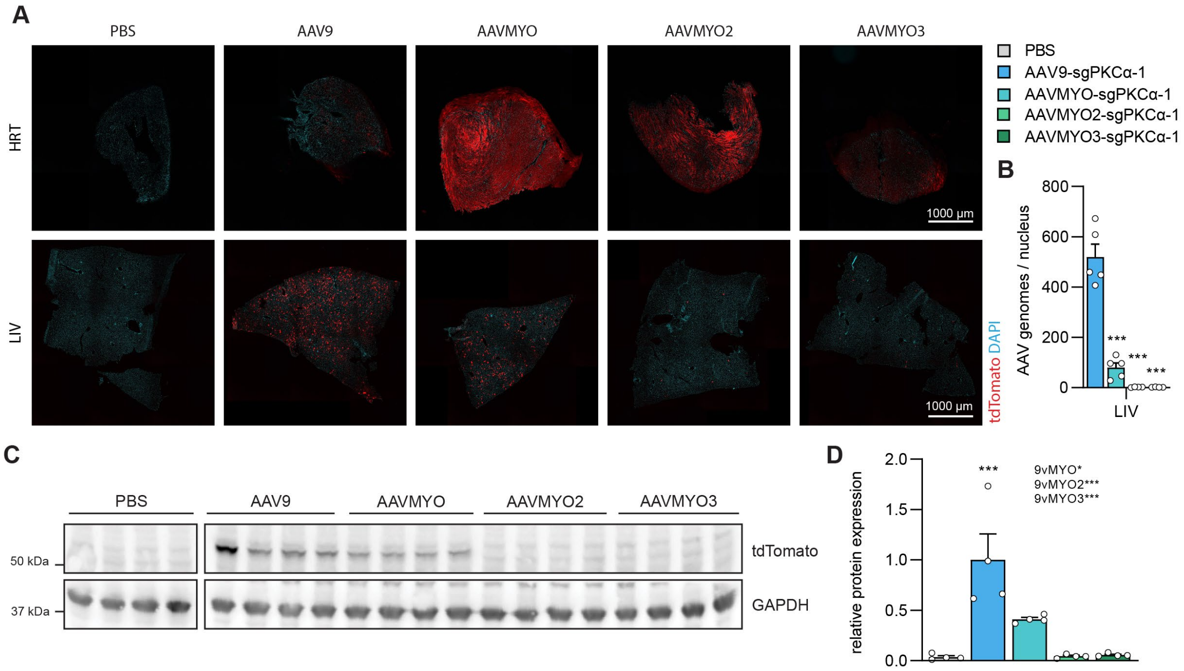


**Figure S7: AAV transduction of heart and liver upon intravenous administration.** (A) Representative images of heart (HRT) or liver (LIV) cross-sections stained for tdTomato (red) and DAPI (blue), 6 weeks post-intravenous injection of PBS or AAV (1.0 x $10^{14}$ vg/kg) into Cas9mKI mice. (B) Distribution of AAVs in liver of Cas9mKI mice upon intravenous injection of PBS (grey), AAV9-sgPKCα-1 (light blue), AAVMYO-sgPKCα-1 (cyan), AAVMYO2-sgPKCα-1 (light green) or AAVMYO3-sgPKCα-1 (dark green). (C) Expression of tdTomato in liver. (D) Quantification of tdTomato expression in liver, normalized to AAV9-sgPKCα-1. Note that AAVMYO2 and AAVMYO3 do not express detectable tdTomato. Data are means ± SEM. N = 4 - 5 mice for each condition. Statistical significance is based on one-way ANOVA with Tukey's post-hoc test. *P < 0.05, **P < 0.01, ***P < 0.001.

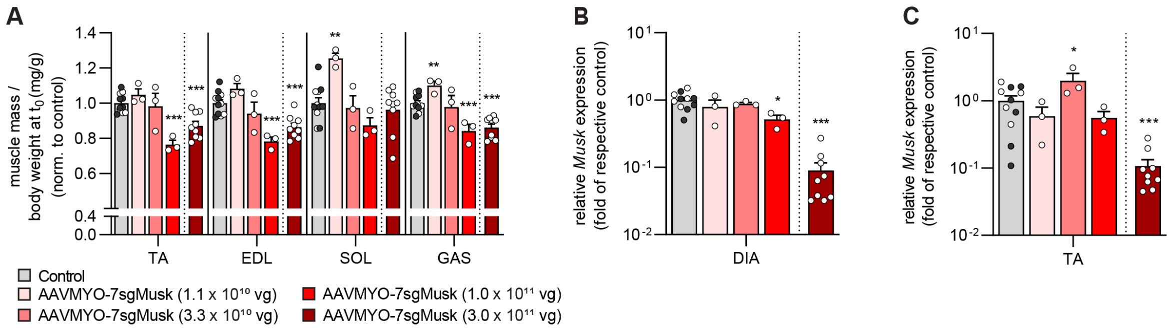


**Figure S8: High doses of AAVMYO-CRISPR/Cas9 induces a systemic knockout of *Musk*.** (A) Changes in mass of contralateral muscles of AAVMYO-7sgMusk-injected mice (red colors) at the indicated dose, compared to control mice (grey bar; white dots indicate PBS-injected Cas9mKI mice; black dots AAVMYO-7sgMusk-injected control mice). (B) Relative mRNA expression of *Musk* in diaphragm and (C) in the contralateral TA muscle of control and AAVMYO-7sgMusk-injected Cas9mKI mice. Data are means ± SEM. N = 3 - 11 mice. Statistical significance is based on unpaired t-test comparing to control. *P < 0.05, **P < 0.01, ***P < 0.001.

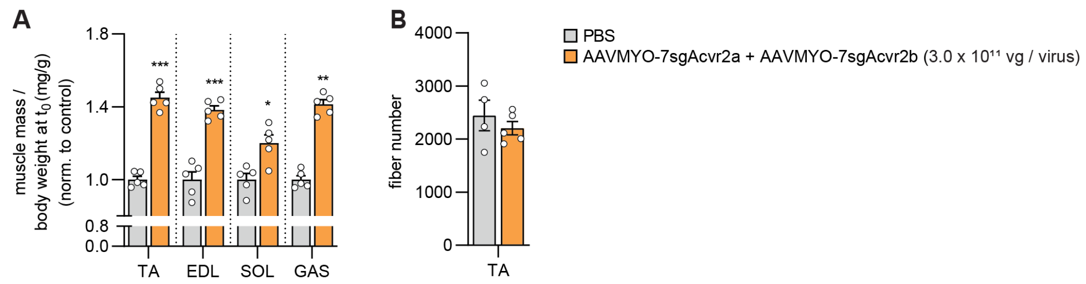


**Figure S9: AAVMYO-CRISPR/Cas9-mediated double knockout of *Acvr2a/Acvr2b* induces systemic hypertrophy without hyperplasia.** (A) Changes in mass of *tibialis anterior* (TA), *extensor digitorum longus* (EDL), *soleus* (SOL) and *gastrocnemius* (GAS) muscles of the contralateral, non-injected leg of AAVMYO-7sgAcvr2a/b- (orange) and PBS-injected mice (grey). (B) Total fiber number of TA muscles injected with PBS (grey) or AAVMYO-7sgAcvr2a/b. Data are means ± SEM. N = 4-5 mice. Statistical significance is based on unpaired t-test. *P < 0.05, **P < 0.01, ***P < 0.001.

| Amplicon | Sample | Total reads | Unmodified reads | Modified reads | Unmodified reads (%) | Modified reads (%) | Insertions | Deletions | Substitutions |
|---|---|---|---|---|---|---|---|---|---|
| sgPKCa | PBS_On-target | 135682 | 132219 | 3463 | 97.4 | 2.6 | 633 | 2275 | 602 |
| | PBS_On-target | 156199 | 146779 | 9420 | 94.0 | 6.0 | 1516 | 7071 | 929 |
| | PBS_On-target | 180456 | 174196 | 6260 | 96.5 | 3.5 | 1044 | 4493 | 785 |
| | PBS_On-target | 147320 | 142053 | 5267 | 96.4 | 3.6 | 858 | 3662 | 862 |
| | AAV9_On-target | 121901 | 100866 | 21035 | 82.7 | 17.3 | 2416 | 18021 | 752 |
| | AAV9_On-target | 122339 | 99979 | 22360 | 81.7 | 18.3 | 3200 | 18354 | 1093 |
| | AAV9_On-target | 90438 | 72908 | 17530 | 80.6 | 19.4 | 2422 | 14678 | 557 |
| | AAV9_On-target | 151031 | 120972 | 30059 | 80.1 | 19.9 | 4694 | 24450 | 1129 |
| | AAVMYO_On-target | 173012 | 128325 | 44687 | 74.2 | 25.8 | 6347 | 37453 | 1349 |
| | AAVMYO_On-target | 143357 | 107893 | 35464 | 75.3 | 24.7 | 5906 | 28743 | 1178 |
| | AAVMYO_On-target | 183297 | 145880 | 37417 | 79.6 | 20.4 | 4853 | 31478 | 1485 |
| | AAVMYO_On-target | 171113 | 134209 | 36904 | 78.4 | 21.6 | 6118 | 29912 | 1145 |
| Off-target1 | PBS_Off-target1 | 162258 | 161007 | 1251 | 99.2 | 0.8 | 0 | 36 | 1215 |
| | PBS_Off-target1 | 233264 | 231445 | 1819 | 99.2 | 0.8 | 1 | 25 | 1793 |
| | PBS_Off-target1 | 223015 | 221178 | 1837 | 99.2 | 0.8 | 0 | 43 | 1796 |
| | PBS_Off-target1 | 185684 | 184046 | 1638 | 99.1 | 0.9 | 0 | 22 | 1616 |
| | AAV9_Off-target1 | 233915 | 231352 | 2563 | 98.9 | 1.1 | 1 | 55 | 2515 |
| | AAV9_Off-target1 | 228564 | 226682 | 1882 | 99.2 | 0.8 | 1 | 29 | 1854 |
| | AAV9_Off-target1 | 272383 | 269940 | 2443 | 99.1 | 0.9 | 2 | 44 | 2399 |
| | AAV9_Off-target1 | 274379 | 272348 | 2031 | 99.3 | 0.7 | 1 | 28 | 2002 |
| | AAVMYO_Off-target1 | 180528 | 178855 | 1673 | 99.1 | 0.9 | 0 | 27 | 1646 |
| | AAVMYO_Off-target1 | 224851 | 222996 | 1855 | 99.2 | 0.8 | 4 | 35 | 1819 |
| | AAVMYO_Off-target1 | 248916 | 247134 | 1782 | 99.3 | 0.7 | 0 | 22 | 1760 |
| | AAVMYO_Off-target1 | 221094 | 218803 | 2291 | 99.0 | 1.0 | 1 | 41 | 2249 |
| Off-target2 | PBS_Off-target2 | 140231 | 139338 | 893 | 99.4 | 0.6 | 0 | 11 | 882 |
| | PBS_Off-target2 | 107946 | 107215 | 731 | 99.3 | 0.7 | 0 | 17 | 714 |
| | PBS_Off-target2 | 163721 | 162511 | 1210 | 99.3 | 0.7 | 0 | 18 | 1192 |
| | PBS_Off-target2 | 167407 | 166321 | 1086 | 99.4 | 0.6 | 0 | 6 | 1080 |
| | AAV9_Off-target2 | 185340 | 183883 | 1457 | 99.2 | 0.8 | 0 | 14 | 1444 |
| | AAV9_Off-target2 | 108346 | 107604 | 742 | 99.3 | 0.7 | 0 | 9 | 733 |
| | AAV9_Off-target2 | 160286 | 159235 | 1051 | 99.3 | 0.7 | 0 | 6 | 1045 |
| | AAV9_Off-target2 | 183724 | 182427 | 1297 | 99.3 | 0.7 | 0 | 9 | 1288 |
| | AAVMYO_Off-target2 | 142479 | 141374 | 1105 | 99.2 | 0.8 | 0 | 9 | 1096 |
| | AAVMYO_Off-target2 | 159580 | 158318 | 1262 | 99.2 | 0.8 | 0 | 13 | 1249 |
| | AAVMYO_Off-target2 | 159491 | 158199 | 1292 | 99.2 | 0.8 | 0 | 10 | 1282 |
| | AAVMYO_Off-target2 | 244032 | 242228 | 1804 | 99.3 | 0.7 | 0 | 11 | 1793 |
| Off-target3 | PBS_Off-target3 | 91217 | 90882 | 335 | 99.6 | 0.4 | 0 | 20 | 315 |
| | PBS_Off-target3 | 119076 | 118513 | 563 | 99.5 | 0.5 | 0 | 38 | 525 |
| | PBS_Off-target3 | 62684 | 62429 | 255 | 99.6 | 0.4 | 0 | 7 | 248 |
| | PBS_Off-target3 | 141915 | 141347 | 568 | 99.6 | 0.4 | 0 | 32 | 536 |
| | AAV9_Off-target3 | 163830 | 163255 | 575 | 99.6 | 0.4 | 0 | 15 | 560 |
| | AAV9_Off-target3 | 65223 | 64837 | 386 | 99.4 | 0.6 | 0 | 6 | 380 |
| | AAV9_Off-target3 | 169763 | 169143 | 620 | 99.6 | 0.4 | 0 | 13 | 607 |
| | AAV9_Off-target3 | 159359 | 158669 | 690 | 99.6 | 0.4 | 0 | 21 | 669 |
| | AAVMYO_Off-target3 | 53238 | 52913 | 325 | 99.4 | 0.6 | 0 | 30 | 295 |
| | AAVMYO_Off-target3 | 85235 | 84916 | 319 | 99.6 | 0.4 | 0 | 9 | 310 |
| | AAVMYO_Off-target3 | 174650 | 173907 | 743 | 99.6 | 0.4 | 0 | 18 | 725 |
| | AAVMYO_Off-target3 | 51100 | 50889 | 211 | 99.6 | 0.4 | 0 | 7 | 204 |
| Off-target4 | PBS_Off-target4 | 131144 | 130499 | 645 | 99.5 | 0.5 | 0 | 10 | 635 |
| | PBS_Off-target4 | 122802 | 122249 | 553 | 99.5 | 0.5 | 0 | 5 | 548 |
| | PBS_Off-target4 | 164119 | 163468 | 651 | 99.6 | 0.4 | 0 | 16 | 635 |
| | PBS_Off-target4 | 111113 | 110709 | 404 | 99.6 | 0.4 | 0 | 1 | 403 |
| | AAV9_Off-target4 | 155484 | 154888 | 596 | 99.6 | 0.4 | 0 | 11 | 585 |
| | AAV9_Off-target4 | 203390 | 202433 | 957 | 99.5 | 0.5 | 0 | 26 | 931 |
| | AAV9_Off-target4 | 178095 | 177445 | 650 | 99.6 | 0.4 | 0 | 10 | 640 |
| | AAV9_Off-target4 | 123382 | 122946 | 436 | 99.6 | 0.4 | 0 | 6 | 430 |
| | AAVMYO_Off-target4 | 152311 | 151674 | 637 | 99.6 | 0.4 | 0 | 5 | 632 |
| | AAVMYO_Off-target4 | 113387 | 112990 | 397 | 99.6 | 0.4 | 0 | 5 | 392 |
| | AAVMYO_Off-target4 | 127566 | 127049 | 517 | 99.6 | 0.4 | 0 | 5 | 512 |
| | AAVMYO_Off-target4 | 156835 | 156281 | 554 | 99.6 | 0.4 | 0 | 36 | 518 |

**Table S1: Amplicon NGS analysis of intramuscular injected TA muscle of Cas9mKI mice**
